# Decoding cryptic diversity of moss populations in a forest-tundra ecotone

**DOI:** 10.1101/2025.06.25.660410

**Authors:** Dennis Alejandro Escolástico-Ortiz, Nicolas Derome, Juan Carlos Villarreal A.

## Abstract

Cryptic speciation is widespread among bryophytes, and it appears common in arctic and subarctic mosses where cryptic lineages may occur sympatrically in the environment. However, cryptic lineages are rarely considered in genetic diversity assessments. This situation poses a challenge, as the complex population structure resulting from cryptic speciation, along with factors like the predominance of clonality, can lead to inaccurate estimates of biodiversity. In this sense, we studied two populations of the subarctic moss *Racomitrium lanuginosum* in the forest-tundra ecotone to test the impact of accounting for differentiated genetic groups on moss genetic diversity estimates, clonal structure and microbial covariation. We performed genotyping-by-sequencing to infer genetic diversity and structure in the forest tundra and the shrub tundra. Genetic groups were identified using haplotype-based coancestry matrices and phylogenetic analyses. The clonal structure was explored by determining multilocus genotypes at the population and a finer scale (225 cm^2^). The covariation between genetic groups and microbial communities (bacterial and diazotrophic) was explored. The recognition of cryptic lineages in genetic diversity estimations revealed differences between habitats that remained undetected when treating *R. lanuginosum* as a single species. Clonal growth seemed to predominate and affect the genetic structure at the local scale. Finally, genetic groups did not host specific microbiomes, suggesting that moss microbial associations in the forest-tundra ecotone did strongly respond to a genetic component. This study highlights the importance of accounting for cryptic lineages in genetic diversity estimations for a precise biodiversity assessment in subarctic and arctic ecosystems.

## INTRODUCTION

Cryptic lineages are genetically differentiated groups with homogeneous characteristics or unclear phenotypic differences, often due to evolutionary processes such as recent divergence, morphological stasis, parallelism, convergence or hybridization (Struck et al., 2018). These cryptic entities can be identified using both morphological and molecular tools (Singhal et al., 2018). Cryptic speciation is widespread in plants (*e.g.*, Les et al., 2015; Liu et al., 2013; Pereira & Prado, 2022; Rees et al., 2023) and animals (*e.g.*, Pfenninger & Schwenk, 2007; Warner, Van Oppen, & Willis, 2015), and appears particularly common in Arctic and Subarctic ecosystems (Brochmann & Brysting, 2008), where species experienced complex demographic histories influenced by climatic oscillations (Abbott et al., 2000; Abbott & Brochmann, 2003; Fedorov et al., 2020; Gargiulo et al., 2019; Hewitt, 2004; Ledent et al., 2019; Westergaard et al., 2019).

Bryophytes, in particular, rely on microscopic characteristics for taxonomic classification making the identification on the field difficult. Bryophytes display cryptic taxa that may arise from genetically isolated populations that were affected by historical and ecological factors while maintaining morphological similarity (Patiño & Vanderpoorten, 2018; Renner, 2020; J. Shaw, 2001). Advances in molecular techniques, including Sanger sequencing, genotyping-by-sequencing, target sequencing, and genome skimming, have successfully identified numerous cryptic bryophyte species (Bechteler et al., 2017; Fuselier et al., 2009; Hedenäs, 2010, 2020a, 2020b; Kyrkjeeide et al., 2016; McDaniel & Shaw, 2003; Medina et al., 2013; Myszczyński et al., 2017; Renner et al., 2018; Ślipiko et al., 2020; Wachowiak et al., 2007; Werner & Guerra, 2004). Notable examples include the peat moss *Sphagnum magellanicum* Brid. (Kyrkjeeide et al., 2016; Shaw et al., 2022), the liverworts *Aneura pinguis* (L.) Dumort. (Wachowiak *et al*., 2007) and *Frullania tamarisci* (L.) (Heinrichs et al., 2010), and the moss *Racomitrium lanuginosum* (Hedw.) Brid. (Hedenäs, 2020a). Notably, *R. lanuginosum* is a widespread dioecious moss inhabiting subarctic, arctic and alpine habitats with a rare presence of sporophytes (a result of sexual reproduction) across its distribution range and it displays predominantly clonal populations (Ellis & Tallis, 2003; Ochyra & Bednarek-Ochyra, 2007; Tallis, 1958). A previous study of Scandinavian populations has shown that *R. lanuginosum* is composed of sympatric molecular lineages or genetic groups with ill-defined morphological traits, making their recognition challenging (Hedenäs, 2020a). Despite significant progress in recognizing cryptic complexes, the classification of these taxa remains controversial due to the lack of standardized diagnostic criteria, sometimes leading to taxonomic inflation and inconsistent taxonomic assignments (*e.g.* cryptic species, sub species, pseudo-cryptic species, varieties, etc.) (Dufresnes et al., 2023).

Assessing genetic diversity in cryptic species may pose a challenge due to the complex spatial genetic structure. Genetic variation in cryptic species may occur between genetic lineages (interspecific) and within populations (intraspecific), creating a nested genetic structure or cryptic diversity (Korshunova et al., 2019). In these cases, interspecific genetic variation arises from cryptic speciation, while intraspecific variation is shaped by population dynamics, with factors such as asexual reproduction further influencing the population genetic structure. For instance, asexual reproduction in bryophytes directly impacts genetic diversity by producing genetically identical individuals (ramets) through structures like gemmae, bulbils, and tubers or by the regeneration of vegetative tissue (*i.e.,* clonality) (Frey & Kürschner, 2011; Newton & Mishler, 1994). Asexual reproduction is common in dioecious bryophytes which produce asexual propagules more frequently than monoecious species (Laenen et al., 2016; Longton, 1992). The production of asexual propagules and clonality shape local genetic structure by influencing the spatial distribution of genets and ramets in the environment (Cronberg et al., 1997; Spagnuolo et al., 2007, 2009; Van Der Velde et al., 2001). Nonetheless, in many situations, predominantly asexual bryophyte populations may exhibit higher genetic diversity than expected. These scenarios are sometimes explained by sporadic sexual reproduction, somatic mutations, and historical population dynamics that contribute to maintaining genetic variation, even in bryophyte clonal systems (Hock et al., 2008; Mishler, 1988; Paasch et al., 2015; Pfeiffer et al., 2006). Incorporating cryptic speciation as a source of genetic variation into analyses allows for a more accurate assessment of genetic diversity and structure in cryptic species (Schuchert, 2014). However, such comprehensive approaches are not yet widely used, often leading to inaccurate estimations of genetic variation in ecosystems.

In addition to morphological traits, some cryptic bryophytes may exhibit distinct ecological characteristics that could aid in their identification. For example, bryophyte lineages are also being characterized through ecological niche analysis (Collart et al., 2021; Hedenäs, 2019; Nieto-Lugilde et al., 2024), providing valuable insights for integrating cryptic species with restricted distributions into conservation strategies (Hedenäs, 2016, 2018). Beyond these ecological assessments, microbial communities associated with bryophytes present a promising yet underexplored avenue (e.g., Rousk, 2022) as a phenotypic extension for characterization of cryptic species. The interaction between bryophytes and their microbiome may be important in local adaptation by influencing the plant ability to respond to environmental stress, for example by enhancing N^2^ acquisition as in the case of the moss-*Nostoc* epiphytic symbiosis (Warshan et al., 2017). Some studies in vascular plants showed that plant genotype influences microbial communities, but its effect is generally weaker than environmental and temporal factors. The genotype signal of microbial communities varies across plant compartments, environments and developmental stages and in some situations, it is only detected through genotype-environment interactions (Brown et al., 2020; Quiza et al., 2023; Semchenko et al., 2021; Wagner et al., 2016). In bryophytes, this knowledge is limited to few studies suggesting that the strength of microbial associations with species or genotypes vary according to the group (Bragina et al., 2012). In peat mosses, cryptic species and genotypes did not directly affect the microbial community composition . It seems that the environment or micro-habitat variation are, in many cases, the key drivers of moss microbial communities. Characterizing bryophyte microbiome could potentially add a distinct trait that help in detecting cryptic species, therefore improving their identification and biodiversity assessments for these often-overlooked groups.

To evaluate how accounting for cryptic species impacts genetic diversity assessments of plants, we focused on two populations of the moss *Racomitrium lanuginosum* in the tundra ecotone, the forest tundra and the shrub tundra in Quebec, Eastern Canada (Fig. 1). A previous studied has demonstrated that *R. lanuginosum* in the forest-tundra ecotone comprises at least two sympatric genetic groups (Escolástico-Ortiz et al., 2023).

**Figure 1.**
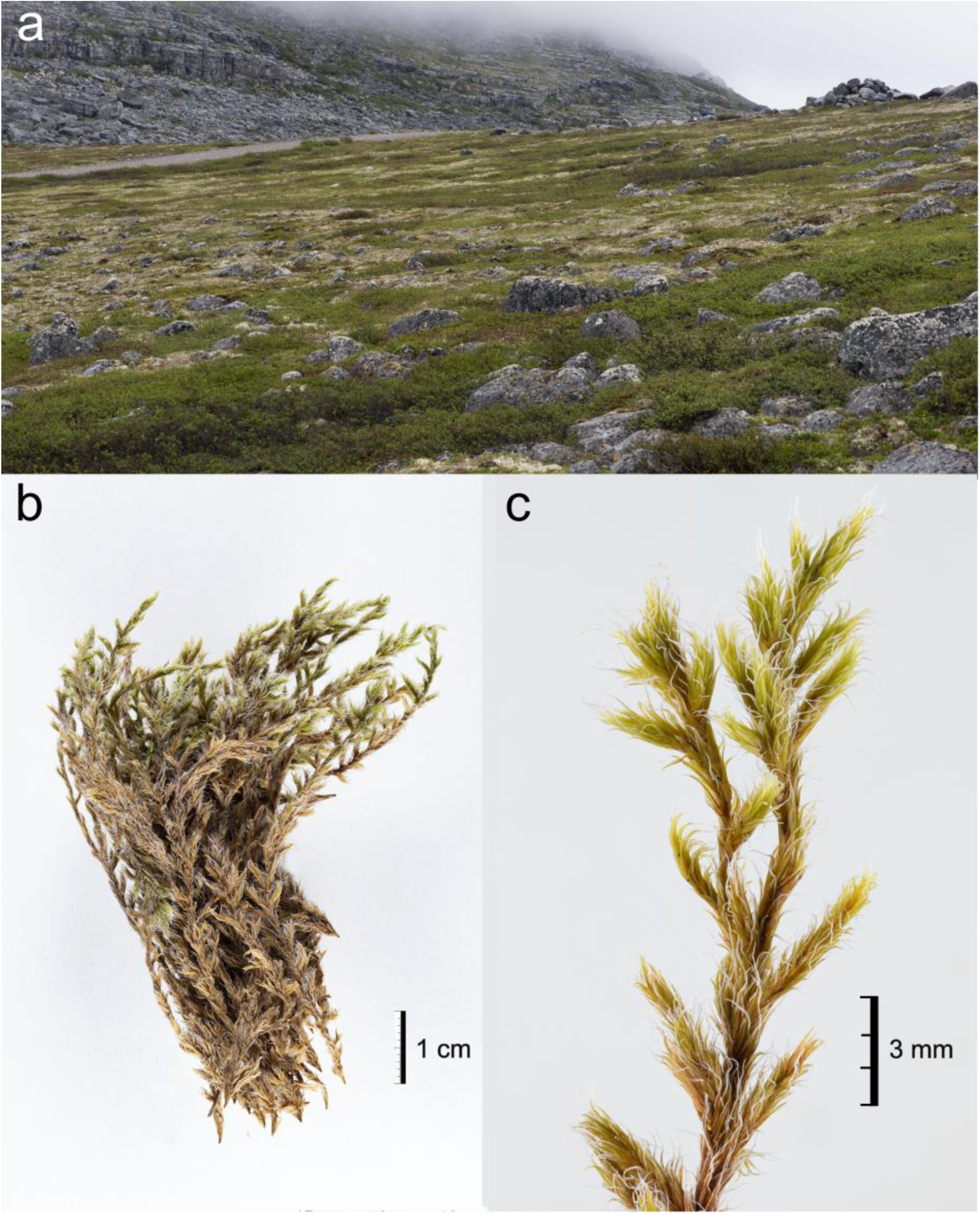
*Racomitrium lanuginosum* habitat and growth form. **a** Whitish mats of *R. lanuginosum* in the shrub tundra at Umiujaq, Canada. **b** Close-up of the moss gametophyte. **c** Details of the shoot (branches) displaying the long hyaline awns. Photos by Catherine Boudreault (a) and Kim Damboise (b,c).

Specifically, we investigated (*i*) how genetic diversity estimates vary whether this moss is treated as one species or two differentiated genetic groups in both habitats, (*ii*) the genetic structure at population and finer-scale (225 cm^2^), and (*iii*) tested whether there is a covariation between the genetic groups and their associated microbial communities (bacterial and diazotrophic groups). We hypothesized that the genetic diversity of populations would vary depending on the number of taxonomical entities considered: a single species or several cryptic species; that genetic variation would be arranged in a nested way with cryptic lineages, genets and ramets; and that genetic groups would host some specific microbial communities. We aimed to demonstrate the importance of carefully decoupling genetic variation to evaluate biodiversity in arctic and subarctic environments.

## MATERIALS AND METHODS

### Study site and sampling design

Sampling was conducted in north-western Quebec near Hudson Bay in June 2019 in two tundra sites near the villages of Kuujjuarapik and Umiujaq (Fig. 2a). Kuujjuarapik is characterized by an ecotone between the boreal forest and the arctic tundra, also known as forest tundra (55°16’30” N; 077°45’30” W). This vegetation type is dominated by small forest patches, shrubs, heaths, lichens and mosses (Payette et al., 2001). The mean annual temperature is -3.6°C, with a mean annual precipitation of 640 mm. The second population corresponds to Umiujaq, a subarctic tundra biome mainly composed of herbaceous species, grasses, lichens and mosses with some scattered low-heigh trees, a mean annual temperature around -4.6°C and mean annual precipitation of 543 mm (Prairie Climate Centre & University of Winnipeg, 2022). The moss populations in this research are referred to as forest tundra for Kuujjuarapik and shrub tundra for Umiujaq.

**Figure 2.**
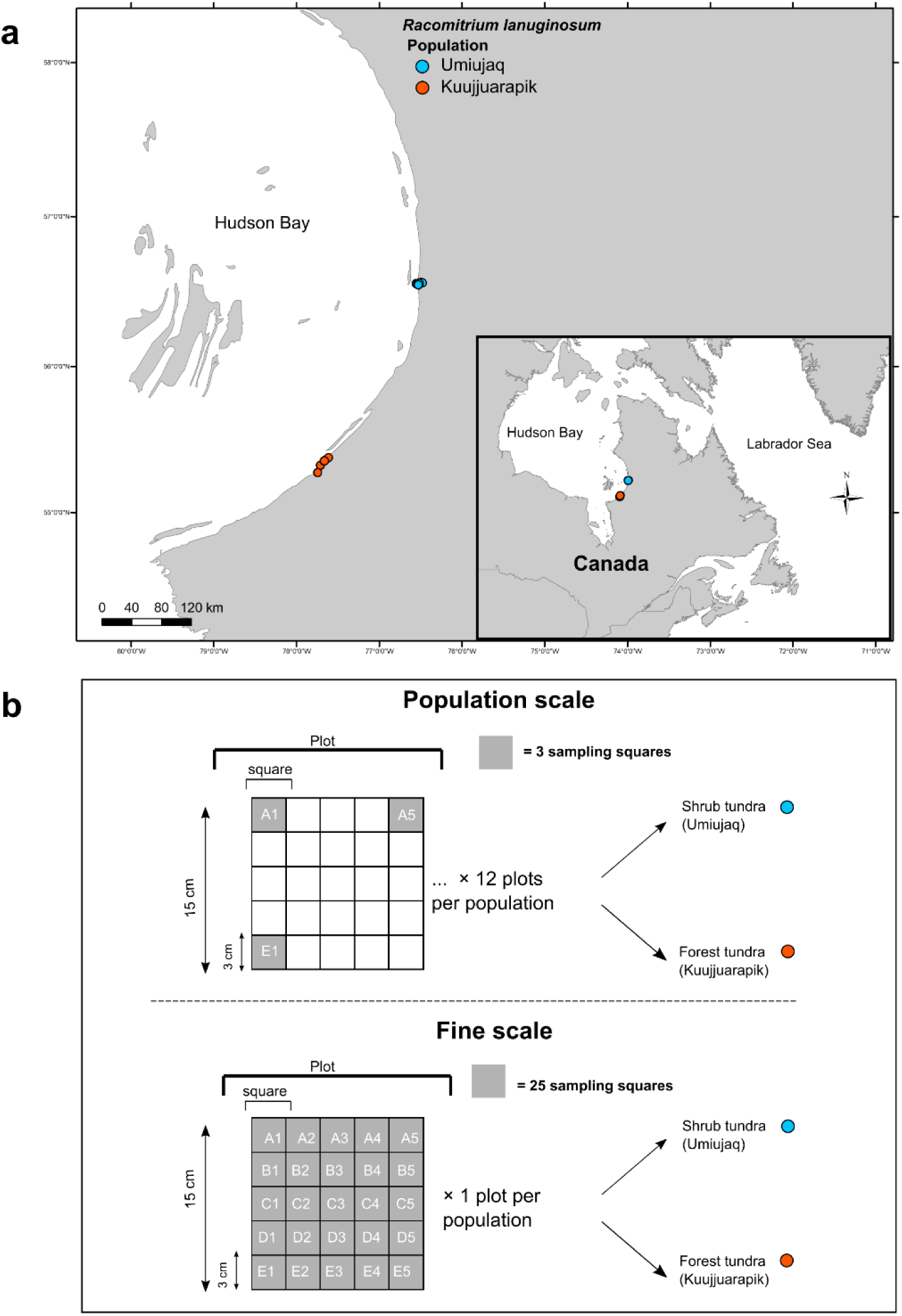
Sampling design of *Racomitrium lanuginosum*’s clonality study. **a** Sampling sites in Eastern Canada. The inset map indicates the location of the two sampling populations in the forest tundra (Kuujjuarapik) and shrub tundra (Umiujaq). **b** Sampling design at population and finer scale levels using 225 cm^2^ plots. At the population scale, we sampled three squares per plot in each population (A1, A5, E1). The fine scale approach comprised all 25 squares of one plot in each population. Measurements are indicated for squares and plots.

The sampling design consisted of 15×15 cm plots with a 3 cm^2^ grid comprising 25 squares coded according to their position (Fig. 2b). Two spatial scales were used to study the genetic structure and diversity. The population scale intended to represent the genetic variation within and among habitats. The sampling for this scale consisted of 14 and 11 plots separated by at least 20 m apart in the forest tundra and shrub tundra, respectively. Only three squares per plot were sampled for this approach, resulting in 33-42 squares per population (75 samples total). We selected three constant squares per plot (A1, A5, E1) to control distance-related genetic variation and collected all the moss shoots within them. The fine-scale approach was designed to describe the clonal structure (genets and ramets) in an area of 225 cm^2^. We used one plot per tundra type and sampled every shoot from each square (25) by putting them in envelopes according to their position code (*e.g.* A1, A2, A3). A total of 50 samples were used for the fine scale. We used one shoot per square for genomic DNA extraction to limit the occurrence of one genet per sample.

Single shoots were homogenized with liquid nitrogen, and genomic DNA was isolated using the CTAB method (Murray & Thompson, 1980). DNA was used for both, GBS to determine the genetic structure and metabarcoding to characterize the microbial community.

### Genotyping-by-sequencing

We used a double-digested (*Pst*I-*Msp*I) genotyping-by-sequencing approach (GBS). Library preparation consisted of 10 µl of DNA with around 20 ng/µl concentrations per sample. Single-end reads were produced using an Ion Torrent technology at the Genomic Analysis Platform of the Institute of Integrative Biology and Systems of Laval University (Québec, Canada). A total of 125 samples were sequenced, of which 75 correspond to the population scale and 50 to the fine scale (see above). The dataset produced 145,105,065 reads.

To confirm the presence of previously recognized cryptic lineages in the study area, we combined our sequences with the data of *R. lanuginosum* from the northern hemisphere and the congeneric *R. geronticum* Müll. Hal. (Escolástico-Ortiz et al., 2023), resulting in 230 samples. Raw reads were checked for quality control using FastQC v.0.11.8 (Andrews, 2010). The Stacks v.2.5 pipeline (Catchen et al., 2013) was used to process reads. First, the Stacks module *process_radtags* was employed to clean, trim to a length of 125 bp and demultiplexed sequences, followed by a second quality check. This step retained 47.20% of the total reads (68,502,829). Next, we performed reference-based alignments against two transcriptomes of two congenerics. We aligned reads to *R. elongatum* Ehrh. ex Frisvoll (ID: ABCD) and *R. varium* (Mitt.) A. Jaeger (ID: RDOO) transcriptomes (Carpenter *et al*., 2019; Leebens-Mack *et al*., 2019) using BWA v.0.7.17 (Li & Durbin, 2009). Then, alignments were transformed into BAM format using Samtools v.1.8 (Li et al., 2009) and input into the *gstacks* module with default settings to construct loci.

### Inferring population genetic structure

We used the *population* module of Stacks for variant calling. Different datasets were produced with varying samples (n) and percentages of shared loci among samples (*-R*) and within samples of the same group (-*r*) (Table S1). To create a “species dataset”, we recovered loci occurring in at least 20% of all samples (-*R* 20) with a haploid condition (*--max-obs-het* 0) and a minimum allele frequency of 0.05 (*--min-maf*) to detect the genetic group membership of all our samples. This dataset consists of 230 samples and 638 SNPs. Similarly, we created a “tundra dataset” with 125 samples corresponding to both tundra habitats and the population and fine scales. We filtered loci presented in at least 80% of samples within populations (*-r* 80) with parameters set as in the first dataset, resulting in 612 SNPs. The vcf files were generated for further analysis. Detailed information on samples is presented in Table S2.

Principal components analyses (PCAs) were performed for each dataset to explore the population structure. The species (n=230; -*R* 20) and tundra (n=125; -*r* 80) datasets were formatted into a *genlight* objects to perform PCAs with the package adegenet v.2.0.2 (Jombart, 2008). The PCAs were plotted using ggplot2 v.3.4.2 package (Wickham, 2016) in R v.4.1.3 (R Core Team, 2017). The PCA of the tundra samples recovered two main clusters represented by a combination of the previously known genetic groups, hereafter genetic groups AB and CD. We performed coancestry matrices for both datasets to estimate the haplotype relationships in both datasets. We formatted the vcf files into haplotype data using Stacks, and constructed coancestry matrices using the nearest neighbour algorithm in fineRADstructure v1.0 (Malinsky et al., 2018).

Additionally, we performed phylogenetic analyses to confirm the genetic membership of the tundra samples using the species dataset. Unlinked SNPs were filtered from the species dataset (n=230; -*R* 20), resulting in 442 variants. We used these SNPs to construct a maximum-likelihood tree in RAxML (Stamatakis, 2014) with the GTR+Γ model and 100 bootstrap replicates, considering the variability of all positions (*--asc-corr* = lewis). The tree was rooted to the congeneric *Racomitrium pruinosum*. A phylogenetic network was also estimated using 2 718 loci from the species dataset in SplitsTree v.4.16.1 (Huson & Bryant, 2006) using the Neighbor Net method and uncorrected p-distances.

### Assessing genetic diversity

To assess the effect of accounting for cryptic lineages, we inferred genetic summary statistics for the tundra dataset. We performed genetic estimates in two scenarios: one considering one single species and another with two genetic groups. First, summary statistics at the population scale (n = 75, samples included in the tundra dataset and sampled at the population scale) were estimated in *populations* of Stacks for each tundra type, treating *R. lanuginosum* as a single species (all genetic groups). To equally compare genetic diversity between tundra types, we corrected the uneven sampling by randomly discarding samples from the forest tundra. The corrected dataset consisted of 33 samples per tundra type. The nucleotide diversity, haplotype diversity and private alleles were estimated for both habitats. Then, we examined population summary statistics using a partition per genetic group and tundra type. The number of samples was corrected as in the previous analyses, resulting in 20 samples for genetic group AB and 46 samples for group CD. Genetic summary statistics were inferred using loci shared by 80% of individuals (*-r* 80) within populations with constant parameters. The haplotype and nucleotide diversity per loci were compared between tundra types to evaluate how the genetic group partitioning (all, AB and CD) influenced the summary statistics. We computed the diversity estimates per loci and checked the data distribution to select the proper statistical test. Due to the non-Gaussian data, Mann-

Whitney tests were used to compare haplotype and nucleotide diversity between tundra types in each dataset. All statistical analyses were performed in R using the package MASS v.7.3-55 (Venables & Ripley, 2002).

### Estimating clonal diversity and structure

Individual genotype assignments were carried out using the package poppr v.2.9.3 (Kamvar et al., 2014). The tundra dataset (n = 125; -*r* 80) was used to determine multilocus genotypes (MLGs) representing genets at different scales. We transformed the vcf file into a *snpclone* object using vcfR v.1.12.0 (Knaus & Grünwald, 2017) and adegenet v.2.0.2 (Jombart, 2008). To assign contracted MLGs, we applied the Hamming distance (i.e., the number of differences between two strings) and the *cutoff_prediction* function, which estimates the threshold for identifying contracted MLGs (Kamvar et al., 2014). The threshold was based on the farthest neighbour algorithm, with a genetic distance of 0.0286 (Fig. S1).

Clonal diversity at the population scale was estimated per genetic group and tundra type. Sample-size corrections were applied to the dataset to allow habitat comparison. The Shannon-Wiener’s H, Stoddart and Taylor’s G, Simpson (lambda) and Evenness indexes were calculated per group using *diversity_stats* in poppr. In addition, we performed analyses of molecular variance (AMOVA) for each genetic group using tundra type and plot as sources of variation. We performed two kinds of AMOVAs, one including all samples (clones included) and the other controlling for the presence of clones. To explore the spatial arrangement of genets and ramets in the fine scale, we identified each sample’s genetic group and MLG within the plot and mapped their spatial location.

The relationship between genetic and geographic distances was investigated through isolation-by-distance (IBD) analyses in dartR v.2.0.4 package (Gruber et al., 2018). We used the 75 samples of the population scale to infer the IBD pattern using genetic (F_ST_) and geographic (log) distances between plots with the function *gl.ibd* and 999 permutations. Additionally, we performed independent genetic group IBD analyses to explore group-specific responses.

### Microbial community analyses

We studied the *R. lanuginosum*’s microbial communities and their co-variation with genetic groups. The detailed methodology, from amplicon sequencing to bioinformatic analyses, is described in Escolástico-Ortiz et al., (2023). Briefly, we amplified the V3-V4 hypervariable regions (347R, 803R) of the *16S SSU rRNA* gene and the *nifH* gene (Ueda19F, R6) to determine the bacterial and diazotrophic community of the moss samples. A two-PCR protocol using the selected primers and Illumina indexes was performed with different PCR programs according to each molecular marker. We purified PCR products, pooled them in an equimolar concentration and sent them for sequencing at the *Plateforme d’analyses génomiques* (IBIS, Université Laval, Quebec City, Canada) using an Illumina MiSeq sequencer (Illumina, San Diego, CA, USA). The sequences were then processed using DADA2 v.1.21 (Callahan et al., 2016) for the 16S rRNA and NIFMAP (Angel et al., 2018) for the *nifH* gene. After quality check and removal of potential contaminants (e.g. chimeric amplicons), the final ASV and OTU tables were produced.

A phyloseq v.1.48 object was created for each molecular marker (*16S SSU rRNA* and *nifH*) by selecting only moss samples with associated genomic information (genetic structure results). All the following analyses were performed for the *16S SSU rRNA* and *nifH* markers independently. Rarefied datasets based on 90% of the minimum sample size (*16S SSU rRNA* = 1073; *nifH* = 1232) were used to infer alpha diversity. We estimated the Shannon and Pielou indexes in vegan v.2.6-6.1 (Oksanen et al., 2018) and the Faith phylogenetic distance in picante v.1.8.2 (Kembel et al., 2010). The effect of habitat and genetic group on alpha diversity (*16S SSU rRNA* and *nifH*) was assessed with linear mixed models (LMM) using lmer4 v.1.1-35.5 package (Bates et al., 2015) in R. We used REML estimation using “habitat” and “genetic group” as fixed effects and “plot” as a random effect to account for non-independence of moss samples. When the variation of the random effect was negligible, we employed linear models (LM) instead. Model assumptions were checked on the DHARMa v.0.4.7 package (Hartig, 2024). Differences between habitats (forest tundra vs shrub tundra) and genetic groups (AB vs CD) were tested with Type III Analysis of Variance (ANOVA) with Satterthwaite’s method for LMM implemented in the lmerTest v3.1-3 package (Kuznetsova et al., 2017) and standard ANOVA for LM.

Microbial taxonomic profiles of moss samples were generated in ggplot2 v.3.5.1 (Wickham, 2016) and are presented per habitat and genetic group. The 20 most abundant taxa are shown in bar plots with the rest pooled as “Other”.

Beta diversity was inferred using normalized datasets with the geometric mean of pairwise ratios (GMPR) method applied in GUniFrac v.1.8 package (Chen et al., 2012). We calculated Bray-Curtis distance matrices and used them as input for non-metric multidimensional scaling (NMDS). A permutational analysis of variance (PERMANOVA) was conducted on vegan package to explore the effects of habitat and genetic group on microbial communities. The heterogeneity of the variance (betadisper) was checked with a permutational test in the vegan package.

Differentially abundant microbial genera between habitats and genetic groups were identified in ANCOM-BC2 v.2.2.2 package (Lin & Peddada, 2024) using the false discovery rate as adjusted p-value method with sensitivity analyses, that allow to identify taxa that are sensible to the zero addition, and default settings (e.g. alpha = 0.05, prv_cut = 0.10, lib_cut = 0).

## RESULTS

### Genetic structure and diversity between habitats

The genetic structure analyses based on 230 samples of *Racomitrium lanuginosum* confirmed the presence of cryptic lineages in the forest tundra and shrub tundra (Fig. 3). However, genetic group memberships were not accurately determined. Instead, the PCA and coancestry matrix of the *R. lanuginosum* tundra samples (n = 125) suggested the presence of two main lineages, AB and CD, related to previously recognized genetic groups mentioned above (Fig. 4 and S2).

**Figure 3.**
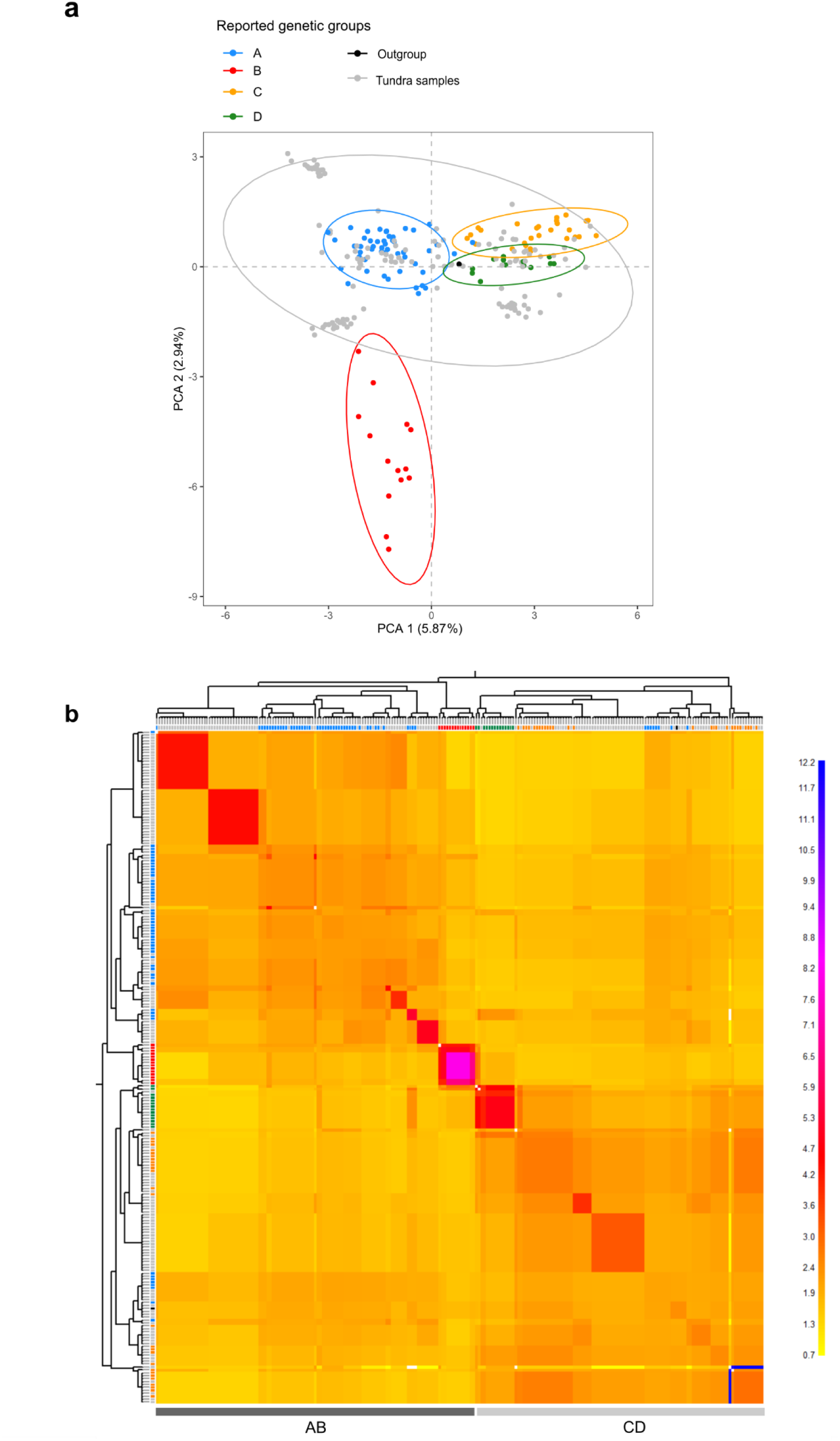
Genetic structure of *Racomitrium lanuginosum* based on 230 samples (-R 20). **a** Principal component analyses based on 638 SNPS. The genetic groups correspond to those previously identified in Escolastico-Ortiz et al. (2023), and the tundra samples are indicated in gray. The dotted circles indicate the 95% confidence intervals for each group. Samples from the tundra are related to two genetic group clusters, AB and CD. **b** Clustered coancestry matrix of haplotypes. The colour scale indicates the level of shared coancestry in terms of loci, ranging from yellow for low to blue for high levels. The left and top axes in the matrix show neighbour trees representing the relationships among samples. Squares at the tree’s tips indicate sample’s genetic group membership: A (blue), B (red), C (orange), D (green), outgroup (black) and tundra samples of this study (gray). Two main groups (AB and CD) were identified, occurring sympatrically in both tundra types. Individuals in the same genetic group tend to share more coancestry within each other than individuals of other groups.

**Figure 4.**
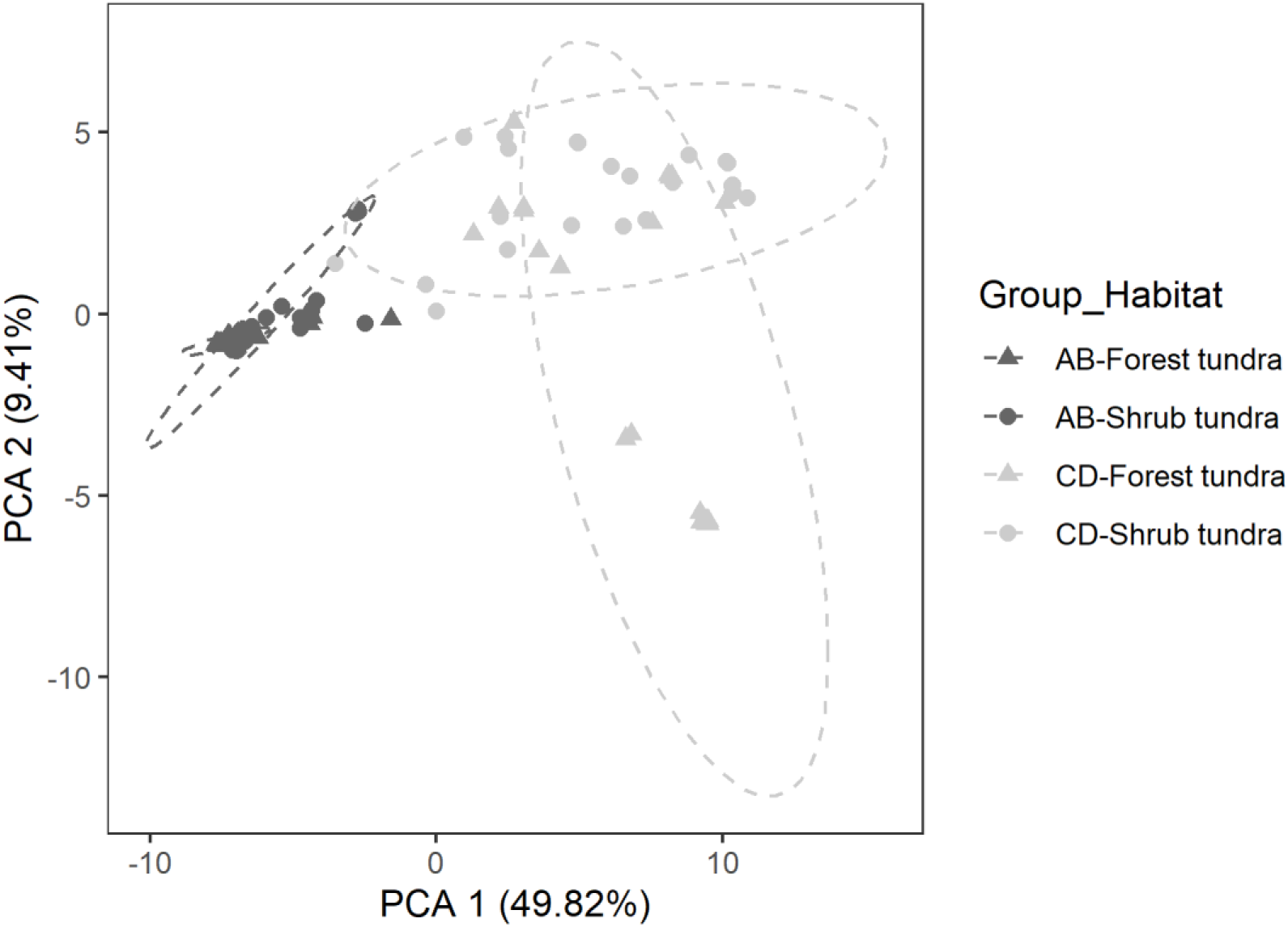
Principal component analyses of *Racomitrium lanuginosum* tundra samples (n=125; -*r* 80) based on 611 SNPs. PCA of SNPs clustered according to the genetic group and tundra type. The dotted circles indicate the 95% confidence intervals for each cluster type. The genetic groups correspond to those found in the genetic structure analyses of the tundra samples. Genetic group membership, not tundra type, better explained the variance of the samples.

Genetic groups occurred sympatrically in both tundra types with no spatial structure and even in the same plot. The samples from the fine-scale approach displayed higher coancestry levels due to the close spatial arrangement (Fig. S2). The maximum likelihood tree and the phylogenetic network of the species dataset (n = 230) did not entirely resolve the relationships among samples but confirmed the close relationships of the fine-scale samples (Figs. S3 and S4).

Genetic summary statistics of *R. lanuginosum* at the population scale with one single specie (no genetic group partitioning) indicated similar genetic diversity in the forest and shrub tundra. However, when genetic groups (AB and CD) were considered separately, estimates of genetic diversity were higher in higher in the shrub tundra compared to the forest tundra for both lineages (Table 1 and Table S3). These group-based estimates suggest that treating genetically distinct lineages as a single taxonomic unit may inflate overall diversity estimates and hide habitat-related genetic differences.

**Table 1.** Genetic summary statistics of *Racomitrium lanuginosum* samples from two tundra types based on genetic groups with corrected sample size. The indexes are estimated for the tundra dataset at the population scale (*r*-80) within habitats and different genetic grouping. The following information is presented: genetic group, haplotype diversity (Hd) and nucleotide diversity (π) per loci with their standard errors (±) for each tundra type, number of samples, U statistic and p-value. The haplotype and nucleotide diversity were inflated when assuming *R. lanuginosum* as a one species (all genetic groups).

| Genetic group | Estimate | Forest tundra | Shrub tundra | Samples | U statistic | p-value |
| --- | --- | --- | --- | --- | --- | --- |
| All | Hd | 0.3673 ± 0.006 | 0.3856 ± 0.006 | 33 | 98,754 | 0.6158 |
| | $\pi$ | 0.3733 ± 0.007 | 0.3922 ± 0.006 | 33 | 96,199 | 0.2464 |
| AB | Hd | 0.247 ± 0.008 | 0.292 ± 0.008 | 10 | 75,817 | 1.81e-05* |
| | $\pi$ | 0.261 ± 0.009 | 0.309 ± 0.009 | 10 | 75,744 | 1.65e-05* |
| CD | Hd | 0.246 ± 0.006 | 0.318 ± 0.005 | 23 | 153,572 | < 2.2e-16* |
| | $\pi$ | 0.251 ± 0.006 | 0.326 ± 0.005 | 23 | 152,993 | < 2.2e-16* |

\* p-value <0.001

### Clonal diversity and spatial structure

Clonal assignments of the *R. lanuginosum* tundra dataset revealed 37 contracted MLGs (i.e. genets) in 125 samples (Table S2 and Fig. S5). The genetic group AB is formed by 15 genets, and group CD is formed by 22. The forest tundra was dominated by a few genets with abundant ramets at the population scale (15 MLGs). The MLG7 and MLG3 were the most abundant genets, representing 12 and 14.67% of all samples, respectively. In contrast, the shrub tundra was composed of several genets (24 MLGs), mainly represented by a single ramet (Table S4). Most of the genets occurred exclusively in one tundra type. Clonal estimates confirmed that the shrub tundra has higher clonal diversity than the forest tundra for both genetic groups (Table 2 and Fig. S6). Molecular variation of both genetic groups was high within plots (Table 3), even after clones were accounted for in the analyses (Table S5).

**Table 2.** Clonal estimates of *Racomitrium lanuginosum* genetic groups in the forest and shrub tundra. The following estimates are presented: number of samples per habitat, number of multilocus genotypes (MLGs), Shannon index (H), Stoddar and Taylor index (G), Simpson index (lambda) and evenness (E.5). The shrub tundra seems to be more diverse than the forest tundra.

| Genetic group | Habitat | Samples | MLGs | H | G | lambda | E.5 |
| --- | --- | --- | --- | --- | --- | --- | --- |
| AB | Forest tundra | 10 | 5 | 1.36 | 3.13 | 0.68 | 0.73 |
|  | Shrub tundra | 10 | 8 | 2.03 | 7.14 | 0.86 | 0.93 |
| CD | Forest tundra | 23 | 8 | 1.64 | 3.60 | 0.72 | 0.62 |
|  | Shrub tundra | 23 | 16 | 2.69 | 13.56 | 0.93 | 0.91 |

**Table 3.** Analyses of molecular variation of *Racomitrium lanuginosum* genetic groups. The variation is analyzed between the forest tundra and the shrub tundra and among plots within habitats or tundra types. The degree of freedom (df), the variances and percentage of the variance explained, the phi-statistics (ɸ), and the p-values are presented. Clones were included in this analysis.

| Genetic group | Source | df | Variance | % variation | Φ-statistics | p-value |
| --- | --- | --- | --- | --- | --- | --- |
| AB | Between habitats | 1 | 3.40 | 6.75 | 0.07 | 0.072 |
|  | Between plots within habitats | 13 | 17.33 | 34.45 | 0.37 | 0.050 |
|  | Within plots | 5 | 29.58 | 58.80 | 0.41 | 0.015* |
|  | Total | 19 | 50.31 | 100 |  |  |
| CD | Between habitats | 1 | 11.93 | 12.84 | 0.13 | 0.001* |
|  | Between plots within habitats | 21 | 15.46 | 16.64 | 0.19 | 0.011* |
|  | Within plots | 23 | 65.49 | 70.51 | 0.29 | 0.001* |
|  | Total | 45 | 92.88 | 100 |  |  |

\* p-value <0.01

Fine-scale clonal analyses indicated that plots were composed of *R. lanuginosum* shoots from different genetic groups and MLGs (Fig. 5). The fine-scale sampling plot in the forest tundra was composed of two genetic groups and four MLGs, with the MLG8 and MLG3 being the most abundant genets for groups AB and CD, respectively. The shrub tundra plot was only represented by group AB and three MLGs, with MLG18 dominating more than half of the plot. The co-occurrence of different genetic groups and genets per plot was also confirmed at the population scale (Table S2). No genets were shared between habitats at the fine scale.

**Figure 5.**
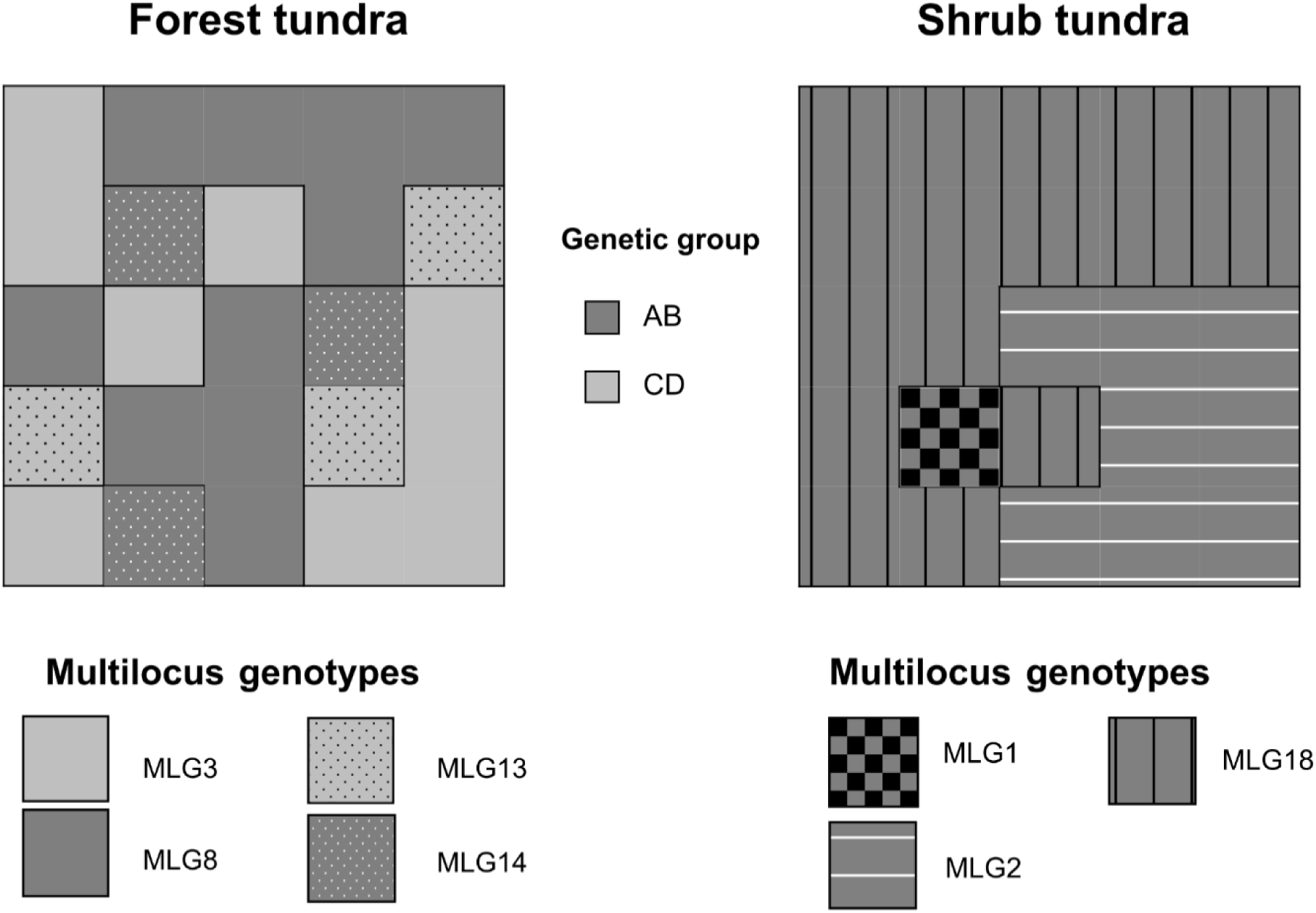
Spatial genetic arrangement of *Racomitrium lanuginosum* shoots in two plots from the tundra. Each square within the plots represents the spatial distribution of shoots. Genetic groups are represented by colors and MLGs (genets) by fill patterns in each habitat. Genetic groups may occur in the same plot as in the forest tundra.

The relationship between genetic and geographic distances (isolation by distance: IBD) of the samples in the tundra habitats (tundra dataset) showed a weak correlation (R^2^ = 0.0129, Mantel R statistic = 0.1138, p-value = 0.011) (Fig. S7a). Genetic group IBD analyses revealed similar results (Fig. S7b-c), showing that the moss can disperse several meters from one plot to another.

### Covariation between genetic groups and microbial communities

The bacterial and diazotrophic communities of *R. lanuginosum* did not covaried with genetic groups. Alpha diversity metrics of bacterial (*16S SSU rRNA*) and diazotrophic (*nifH*) communities suggested that there are no differences between genetic groups (Tables S6-S10). The microbial taxonomic profiles showed little variation between habitats and similar microbial relative abundance between genetic groups at the phylum (Fig. S9) and genus level (Fig. 6a and 6c; Fig. S10). Microbial beta diversity was not significantly affected by *R. lanuginosum* genetic groups according to NMDS ordination and PERMANOVA analyses (Figure 6b and 6d; Table S11). Finally, differential abundant analyses of the *16S SSU rRNA* data recovered 21 potential bacterial genera (from which 10 were unassigned) that may be related to genetic groups.

**Figure 6.**
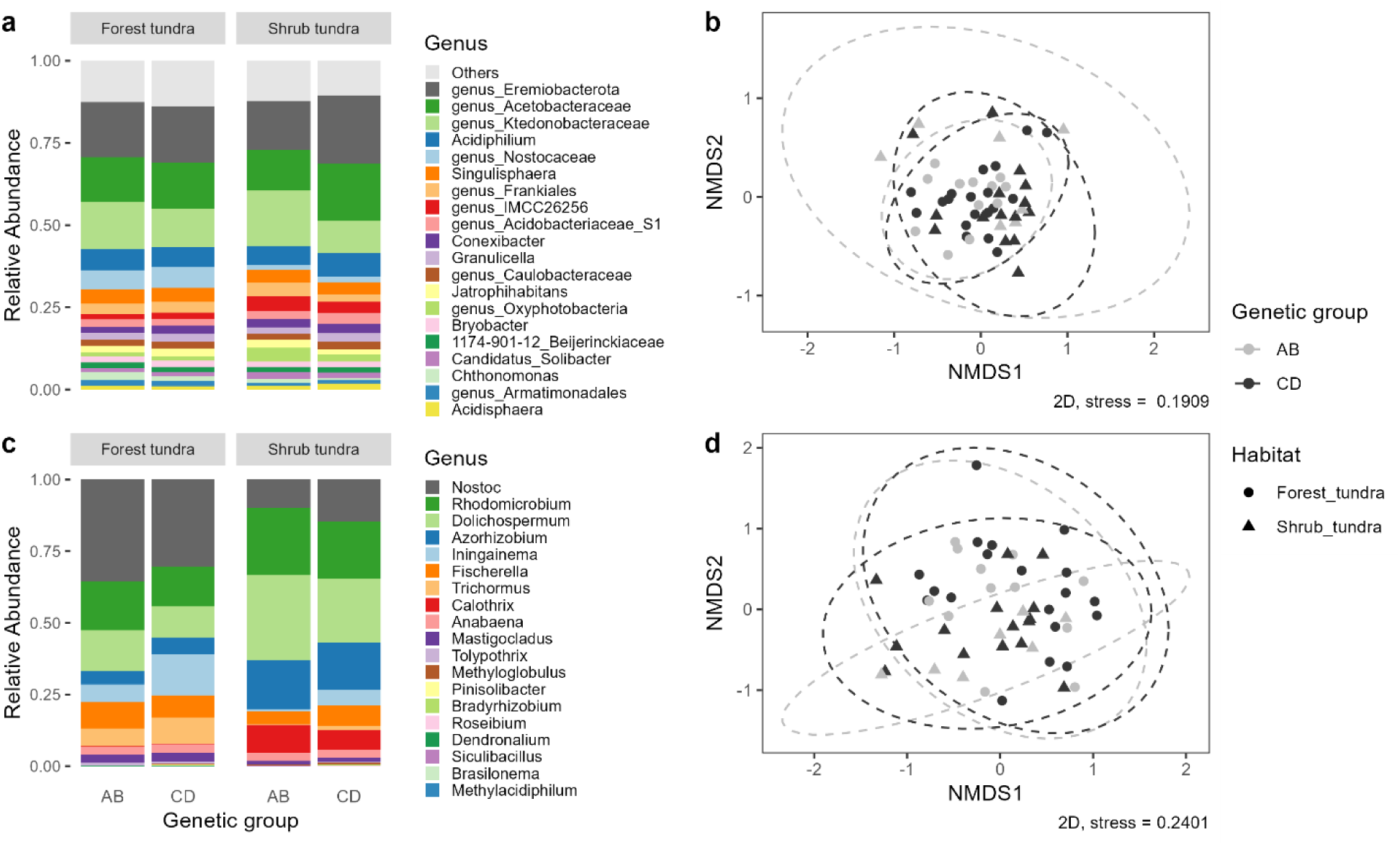
Microbial community profiles and ordinations of microbial communities of *Racomitrium lanuginosum*. **a** Relative abundance of bacterial genera based on the *16S SSU rRNA* gene with samples pooled according to genetic group and habitat. **b** Non-metric multidimensional scaling (NMDS) based on Bray-Curtis distances of *16S SSU rRNA* data. Genetic groups and habitat are represented by colors and shapes, respectively. **c** Relative abundance of diazotrophic genera based on the *nifH* gene with samples pooled according to genetic group and habitat. **d** Non-metric multidimensional scaling (NMDS) based on Bray-Curtis distances of *nifH* data. Genetic groups and habitat are represented by colors and shapes, respectively. The microbial community seem to be less affected by the genetic groups than the habitat type.s

Nonetheless, only two unassigned genera from Fimbriimonadaceae and Hypomicrobiales (Rhizobiales) successfully passed the sensitive analyses of ANCOMBC-2, which control for false positives taxa in pseudo-count transformations (Fig. S11a-b). Differential abundant analysis for the *nifH* data only recovered the genus *Pinisolibacter* in the genetic group CD that did not pass the sensitivity test (Fig. S11c). According to our results, the genetic groups of *R. lanuginosum* did not seem to host specific bacterial (*16S SSU rRNA*) or diazotrophic (*nifH*) communities.

## DISCUSSION

### Recognition of cryptic species uncovers genetic diversity differences between habitats

We explicitly tested how plant genetic diversity was affected by the recognition of cryptic molecular lineages. Populations of the moss *Racomitrium lanuginosum* in the forest tundra and the shrub tundra comprised sympatric genetic groups. Genetic groups corresponded to previously reported cryptic molecular lineages of *R lanuginosum* across its geographic distribution (Escolástico-Ortiz et al., 2023). These genetic groups occurred mixed in the same tundra type and even in the same plot, with no apparent spatial segregation. The presence of cryptic lineages in *R. lanuginosum* has also been highlighted in the Scandinavian peninsula (Hedenäs, 2020a). Even though *R. lanuginosum* genetic groups seem to be distributed in areas with different mean temperatures, potentially suggesting environmental adaptation, evidence of spatial distribution models of molecular lineages from Scandinavia showed no significant differences in the distribution range among groups (Collart et al., 2021). Other moss cryptic lineages also show overlapping distribution patterns, such as *Hamatocaulis vernicosus* (Mitt.) Hedenäs cryptic taxa in southern Sweden (Hedenäs, 2018), *Sphagnum magellanicum* complex in the Northern Hemisphere (Kyrkjeeide et al., 2016) and *Lewinskya affinis* (Brid.) F. Lara, Garilleti & Goffinet cryptic species (Vigalondo et al., 2019). As for the species mentioned above, this pattern seemed to be shaped by the isolation of cryptic lineages in different refugia during Pleistocene glaciations and their subsequent dispersal into new regions (Désamoré et al., 2016; Hedenäs, 2014; Ledent et al., 2019). Sympatric cryptic speciation has been reported in other plants, such as the aquatic herb *Najas canadensis* Michx. due to hybridization events (Les et al., 2015). In the case of *R. lanuginosum,* the cryptic lineages seem to result from isolation of refugial populations, glacial bottlenecks, and long-distance dispersal. Nonetheless, an extended sampling and more explicit analyses, such as Bayesian population modelling, are required to support the processes that contribute to the differentiation of *R. lanuginosum* cryptic taxa.

Our findings support our first hypothesis that treating a species cryptic complex as a single entity impacts the accuracy of across-habitat genetic estimates. The comparisons of genetic diversity with and without genetic grouping proved that habitat genetic variation can be concealed when pooling cryptic lineages into one species. Notably, genetic diversity in the shrub tundra was higher for AB and CD lineages, but no difference was detected between habitats when assuming a single species for *R. lanuginosum*. Differences in genetic composition across habitats were also detected in Scandinavian populations of five mosses, including *R. lanuginosum*, probably due to differences in colonization sources or ecological adaptation in the northern and mountain ranges of Scandinavia (Hedenäs, 2019). Masked genetic diversity is not unique to plants and has also been uncovered in other predominantly clonal species complexes through molecular techniques, highlighting how clonality can obscure underlying genetic structure. In the coral *Seriatopora hystrix* Dana, distinct genetic groups occur sympatrically in the Great Barrier Reef, but independent analyses discerning genetic entities allowed the identification of connectivity patterns and spatial genetic structure within groups (Warner et al., 2015). Similarly, hidden genetic variation was revealed in the hydroid *Plumularia setacea* (L.) with mitochondrial (*COI*) and nuclear markers (*16S SSU rRNA*, *ITS*) across its distribution range; this variation was arranged in several sibling species with fine geographic structure and high genetic diversity within species (Schuchert, 2014). We demonstrated that the concept of *R. lanuginosum* as a unique taxonomic entity inflated the genetic estimates per habitat and concealed the higher genetic variation in the shrub tundra. Therefore, it is essential to account for well-defined molecular lineages in species complexes when conducting genetic assessments to ensure the most accurate and reliable conclusions. Failing those lineages has significant conservation implications, as it can lead to inaccurate biodiversity assessments in protected areas (Feckler et al., 2014). By dissecting genetic variation into cryptic lineages, genotypes and clones (genets and ramets), we can better understand cryptic diversity in environments previously thought to have low evolutionary potential, like the Arctic and Subarctic.

### Clonality influences genetic variation within habitats

Populations of *R. lanuginosum* inhabiting the forest and shrub tundra are characterized by clonal growth and rare sexual reproduction. No sporophytes were recorded in either habitat, which is consistent with findings from Japan (Korpelainen et al., 2012) and the Mediterranean and Eastern continental Europe (Tallis, 1959). The apparent absence of sporophytes in the studied populations could be related to the species phenology. A previous report in a short-growth season habitat indicated that antheridia (male gamete) of *R. lanuginosum* reach maturity after 7-10 months with an inactive phase under the snow layers from mid-December to late April (∼4 months) (Maruo & Imura, 2016). These observations suggest that fertilization may take even a year, assuming that most antheridia mature simultaneously, and as a result, sporophytes would develop in the next growing season. Because our sampling was conducted in July, we could hypothesize that most male gametes were immature for fertilization at that moment and, consequently, no sporophytes were recorded. Additionally, even if sporophytes and spores are produced, the chance of establishment by spores in *R. lanuginosum* seems low compared to clonal growth. Early experimental cultures of the species indicated that once the spore germinated (19-59% after six days), the protonema’s growth rate is low (69.7% after a month), with no bud formation and susceptible to desiccation (Tallis, 1959). It seems that for *R. lanuginosum*, clonal growth represents a suitable strategy to cope with extreme environmental conditions in arctic and subarctic habitats.

The clonal analyses of *R. lanuginosum* validated our second hypothesis regarding the occurrence of the spatial structure of clones at the population and plot levels. Genets seemed to be spatially restricted to one tundra type and do not occur in both habitats, which implies that moss populations in the Canadian tundra persist and disperse locally via clonal growth. This is similar to the genetic partition observed in species of the *Sphagnum magellanicum* complex (Shaw et al., 2023). In addition, most genetic variation of *R. lanuginosum* was found within plots, with different genetic groups and genets intermingled. The observed clonal distribution pattern is typical of the “guerilla” growth form, where distinct genets grow amalgamated in the same patch in a condensed manner (Doust, 1981). A guerilla-like growth form is also common in other northern moss species, such as *Hylocomium splendens* (Hedw.) Schimp. (Cronberg, 2002) and some *Sphagnum* spp. (Shaw et al., 2023). Different genets occurring at close distances increase the likelihood of recombination and reduce self-fertilization and this is particularly important for maintaining genetic diversity in dioecious species with clonal populations where gametes occur on physically separated plants (Johnson & Shaw, 2015). The *R lanuginosum* fine clonal structure and the IBD results indicate that clonal propagation is limited to a several meters in the same habitat. For *R. lanuginosum*, the intermingled arrangement of genets might explain the high source of variation at the plot scale and corroborates the importance of clonal growth for maintaining the existing genetic pool in dioecious bryophytes.

We found higher clonal variation in the shrub tundra than in the forest tundra, with more genets represented by few ramets in the first habitat. It is not unusual for bryophyte populations with rare sexual reproduction and dominance of clonal growth to display more genetic variation than expected for this scenario. Clonality could also be a way of preserving locally adapted genets or rare ones for facing environmental change. For example, in Mediterranean populations of the moss *Pleurochaete squarrosa* (Brid.) Lindb., the predominance of clonal propagation was confirmed, yet high gene diversity (0.644 to 1.000) was detected using inter-simple sequence repeat molecular markers (Spagnuolo et al., 2007, 2009). Similarly, genetic diversity (AFLP fingerprinting) of the moss *Syntrichia caninervis* Mitt. in the Mojave Desert did not differ between sex-expressing and non-sex-expressing populations (Paasch et al., 2015). Recently, genetic diversity in the clonal hornwort *Nothoceros aenigmaticus* J. C. Villarreal and K. D. McFarland was assessed using SNPs across the Appalachians and Mexico indicating varying levels of diversity across populations (Alonso-García et al., 2020). Genetic variation in bryophyte populations with a dominance of clonal growth seems to be mainly driven by the occasional production of sporophytes, complex historical demography, and/or potential somatic mutations (Alonso-García et al., 2020; Kyrkjeeide et al., 2014; Shaw et al., 2023; Spagnuolo et al., 2009). In the case of the fores-tundra ecotone, the differences in genetic diversity may also be related to competition pressures. In the forest tundra, trees can establish and grow faster than *R. lanuginosum*, reducing the available substrate for the establishment and, consequently, the number of potential migrants. In contrast, the relatively bare rock surface in the shrub tundra increases the probability of moss colonization. The impact of vascular plants on the establishment of *R. lanuginosum* has already been mentioned for British populations, indicating that factors such as shading effect, litter accumulation and faster growth rates can interfere with the moss establishment (Tallis, 1959). The forest-tundra in northern Quebec is susceptible to climate change and the responses of trees and shrubs suggest changes in growth patterns that can alter the plant community composition (Labrecque-Foy et al., 2023), potentially affecting the population dynamics of *R. lanuginosum* in the future. All the factors mentioned above coerce to render clonality a suitable means of persistence for a plant with cryptic molecular lineages in the subarctic.

### No significant microbial covariation with genetic groups in *Racomitrium lanuginosum*

The third hypothesis of the research was tested and indicate that Genetic groups in *Racomitrium lanuginosum* did exhibited a strong signal for genetic groups and distinct microbial communities (bacterial or diazotrophic). There were no significant differences in terms of microbial alpha diversity, beta diversity or differentially abundant microbial genera. These results suggest that microbial associations in *R*. *lanuginosum* in the forest-tundra ecotone are primarily influenced by other factors, such as the environment or micro-habitat, rather than genetic differences. Most of the recovered differential abundant bacteria were unassigned taxa, making the inference of moss-microbial interactions difficult to interpret, however, the presence of bacteria from the Rhizobiales in *R. lanuginosum* genetic group CD, suggest potential benefits for the broader plant community. Of note, some bacteria strains in Nocardioides (genetic group CD) have been reported to shown antimicrobial activity (Gesheva & Vasileva-Tonkova, 2012). Our results agree with findings from the *Sphagnum magellanicum* complex, where no significant genetic signals were detected to influence the microbiome of cryptic species, genets, or moss sexes (Shaw et al., 2023). The only observed difference in microbial richness was found at a single sampling location between plant sexes (*Sphagnum divinum* in St. Regis Lake). Similarly, bacterial communities of two morphologically distinct species, *S. fallax* and *S. angustifolium*, were similar (Bragina et al., 2012). A comparable study on reindeer lichens in Quebec revealed that bacterial communities of *Cladonia* species did not exhibit microbial selectivity when considering molecular lineages (Alonso-García et al., 2020), although some differences were observed between species with distinct morphological features. Overall, the microbiome of *R. lanuginosum* genetic groups was not a key trait for cryptic species identification. However, exploring ecological traits to characterize cryptic species remains valuable, particularly when morphological features are homogeneous or inconspicuous.

## CONCLUSION

The subarctic populations of the moss *R. lanuginosum* displayed genetic variation structured in cryptic molecular lineages, revealing the complexity of molecular variation patterns. Accurately recognizing these lineages was crucial for estimating the diversity of two tundra populations. This approach is especially relevant for plant genetics, as the inaccurate circumscription of cryptic taxa can lead to concealed genetic variation and inexact diversity estimates across habitats. Moreover, our results demonstrate that populations of the moss *R. lanuginosum* in the Canadian tundra persist mainly by clonal growth. Clonality influenced the local genetic variation by promoting the spread of dominant clones within habitats. The lack of sporophytes in the populations could be linked to the moss phenology and harsh environmental conditions in the Subarctic. Genetic groups in this species did not host specific microbial communities suggesting that moss microbial associations in the forest-tundra ecotone did strongly respond to a genetic component. *Racomitrium lanuginosum* is a vital element of plant diversity in northern ecosystems and serves as an excellent example of how evolutionary and biological processes, such as cryptic speciation and clonality, can conceal molecular variation across landscapes. This research demonstrates the importance of determining genetic partition in complex biological entities in the Subarctic/Arctic for biodiversity assessments. In the long term, this approach could be useful to identify priority areas for conservation management in terms of genetic diversity in one of the most vulnerable regions on the planet.

## ACKNOWLEDGEMENTS

DAEO thanks the International Association of Bryologists (IAB) for its financial help through the Stanley Greene Award and the “Fonds de recherche du Québec –Nature et Technologies” through the Merit scholarship program for foreign students (PBEEE: https://doi.org/10.69777/307761) and CONACYT for a partial doctoral fellowship. We recognize the support of the Herbier Louis-Marie (QFA). We would also like to sincerely thank Kim Damboise and Catherine Boudreault for their invaluable help and support during fieldwork. We also acknowledge the facilities provided by the Centre of Northern Studies (CEN in French) at Kuujjuarapik and Umiujaq. This research was financed by the Discovery Grant (NSERC): RGPIN-2016-05967 and the Canadian Foundation for Innovation (CFI): 39135. This article is part of the doctoral thesis of DAEO.

## DATA AVAILABILITY

Voucher specimens of each sample were deposited in QFA herbarium and assigned a unique code (see Table S2). Sequences generated in this research are stored in the Sequence Read Archive repository (NCBI) under the BioProject PRJNA1023519. The sequence accession numbers for the samples include SRR26265052 to SRR26265159. Previously published sequences can be retrieved from the BioProject PRJNA735773. Bioinformatic scripts are available on: https://github.com/escolastico-ortizda.

## Notes

### Competing Interest Statement

The authors have declared no competing interest.

