## Supplementary Material - Figures for "Decoding cryptic diversity of moss populations in a forest-tundra ecotone"

### Journal

Dennis Alejandro Escolástico-Ortiz[1,2,3,*], Nicolas Derome[1,2] & Juan Carlos Villarreal[1,2,3]

[1]Département de Biologie, Université Laval, Québec, G1V 0A6, Canada

[2]Institut de Biologie Intégrative et des Systèmes (IBIS), Université Laval, Québec, G1V 0A6, Canada.

[3]Centre d’études nordiques (CEN), Université Laval, Québec, QC, Canada

* ****

### Supplementary Information

### Figures

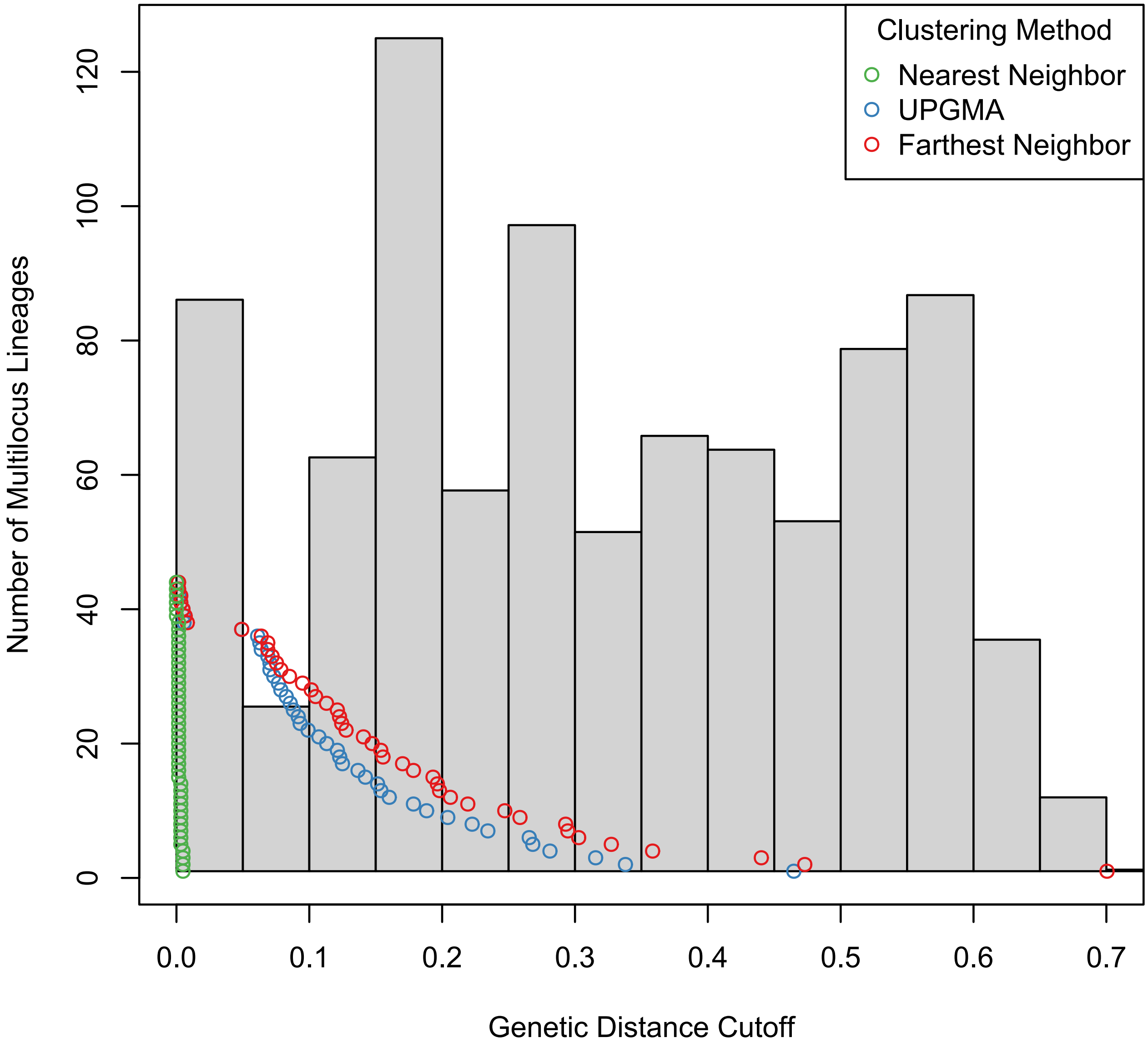


**Figure S1** Genetic distance cutoff in response to the number of *Racomitrium lanuginosum* multilocus genotypes using three clustering methods. The UPGMA and the farthest neighbour gave consistent results.

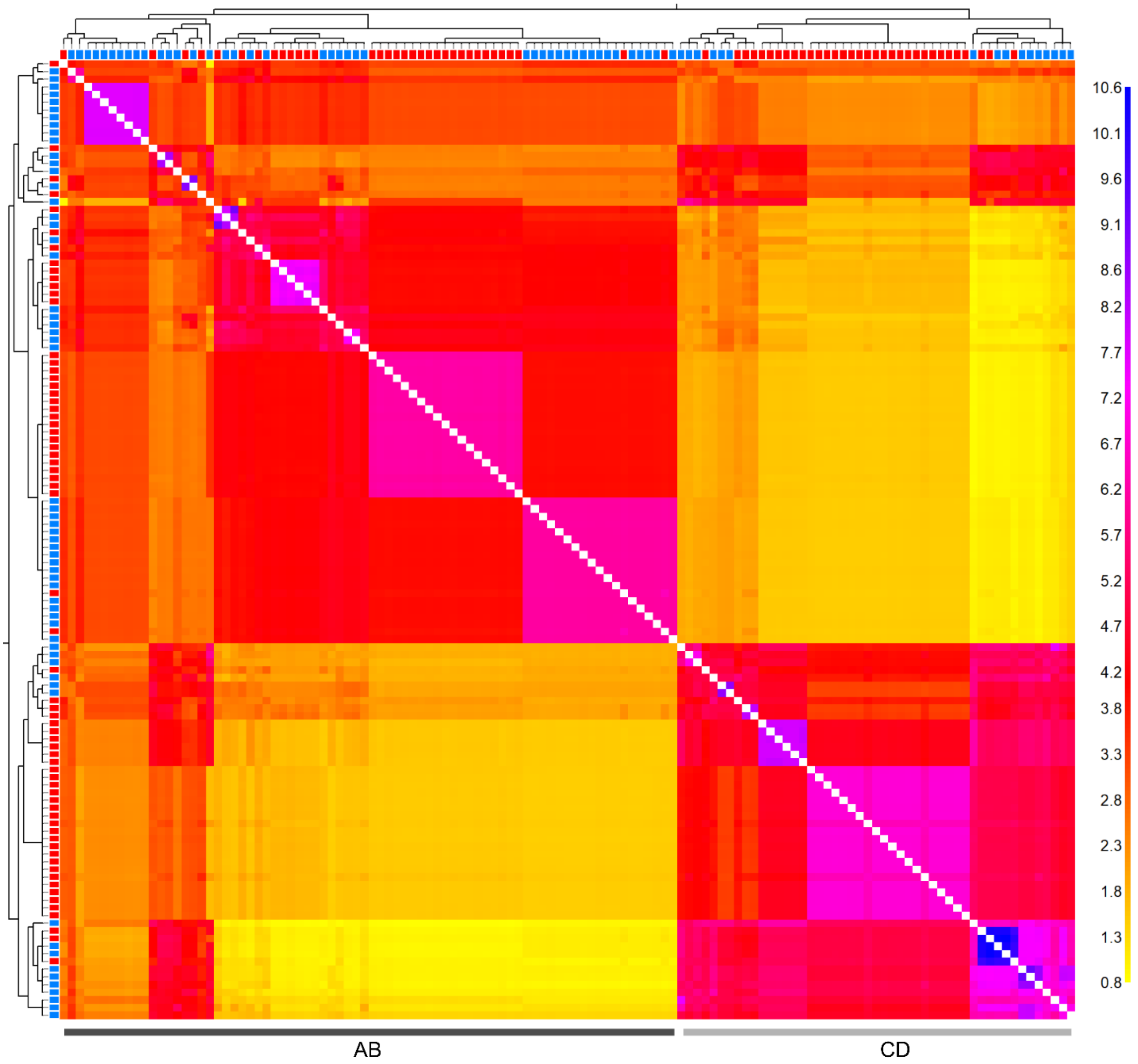


**Figure S2** Coancestry matrix of *Racomitrium lanuginosum* based on haplotypes (611 SNPs) shared by at least 80% of 125 samples (tundra dataset). On the right, the colour scale indicates the level of shared coancestry in terms of loci, ranging from yellow for low to blue for high levels. The left and top axes in the matrix show neighbour trees representing the relationships among samples. Squares at the tree's tips indicate each sample's origin, with red representing the forest tundra and blue the shrub tundra. Two genetic groups (AB and CD) were identified according to shared coancestry, occurring sympatrically in both tundra types. Individuals in the same genetic group tend to share more coancestry within each other than among individuals of other groups.

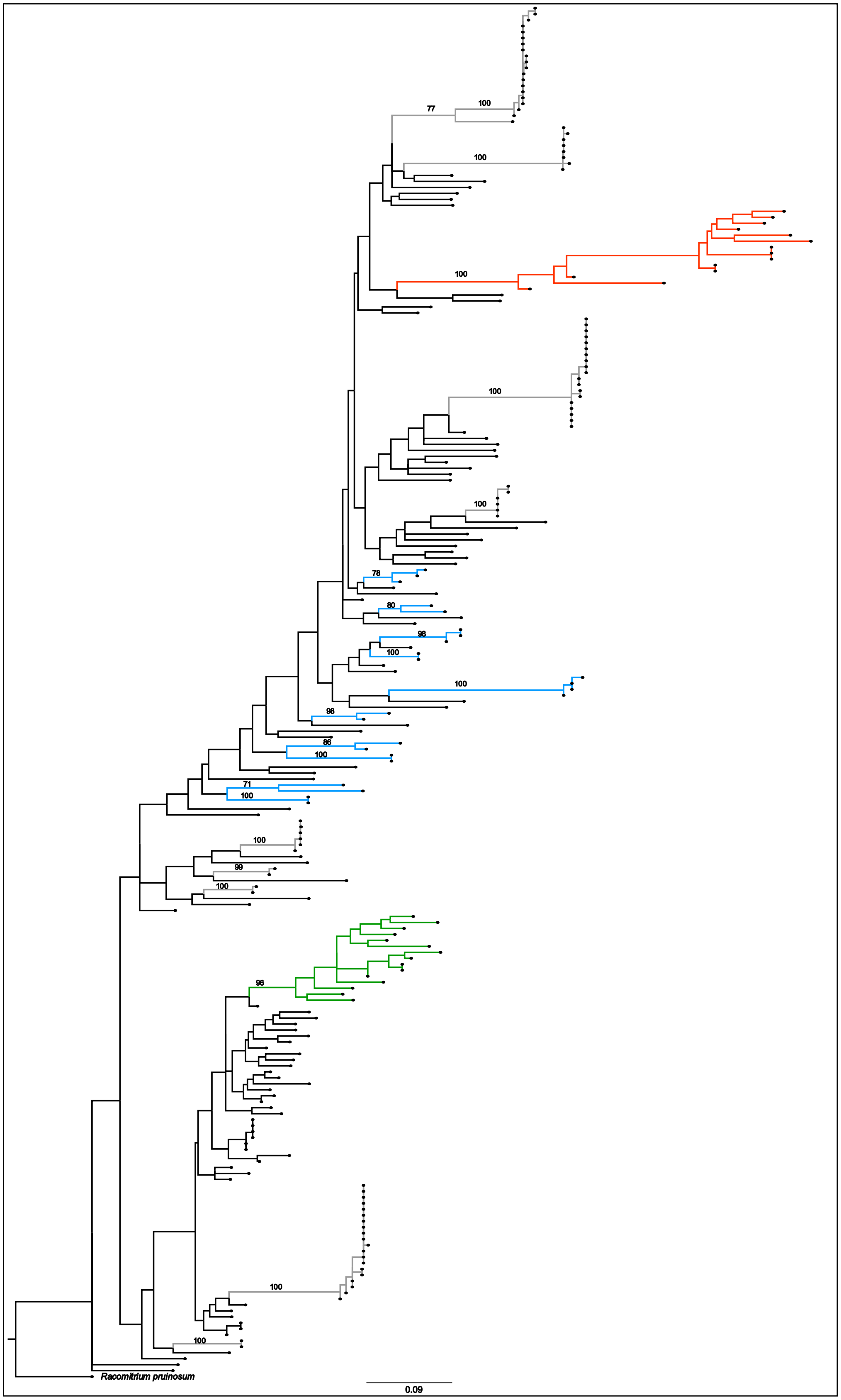


**Figure S3** Maximum likelihood tree of *Racomitrium lanuginosum* samples using *R. pruinosum* as an outgroup (species dataset). The inferred tree is based on 442 unlinked SNPs found in at least 20% of 230 samples (species dataset). Only clades with bootstrap values

>70 are shown and represented in colors according to genetic groups A (blue), B (red) and D (green). Clades represented by the tundra samples are indicated in gray.

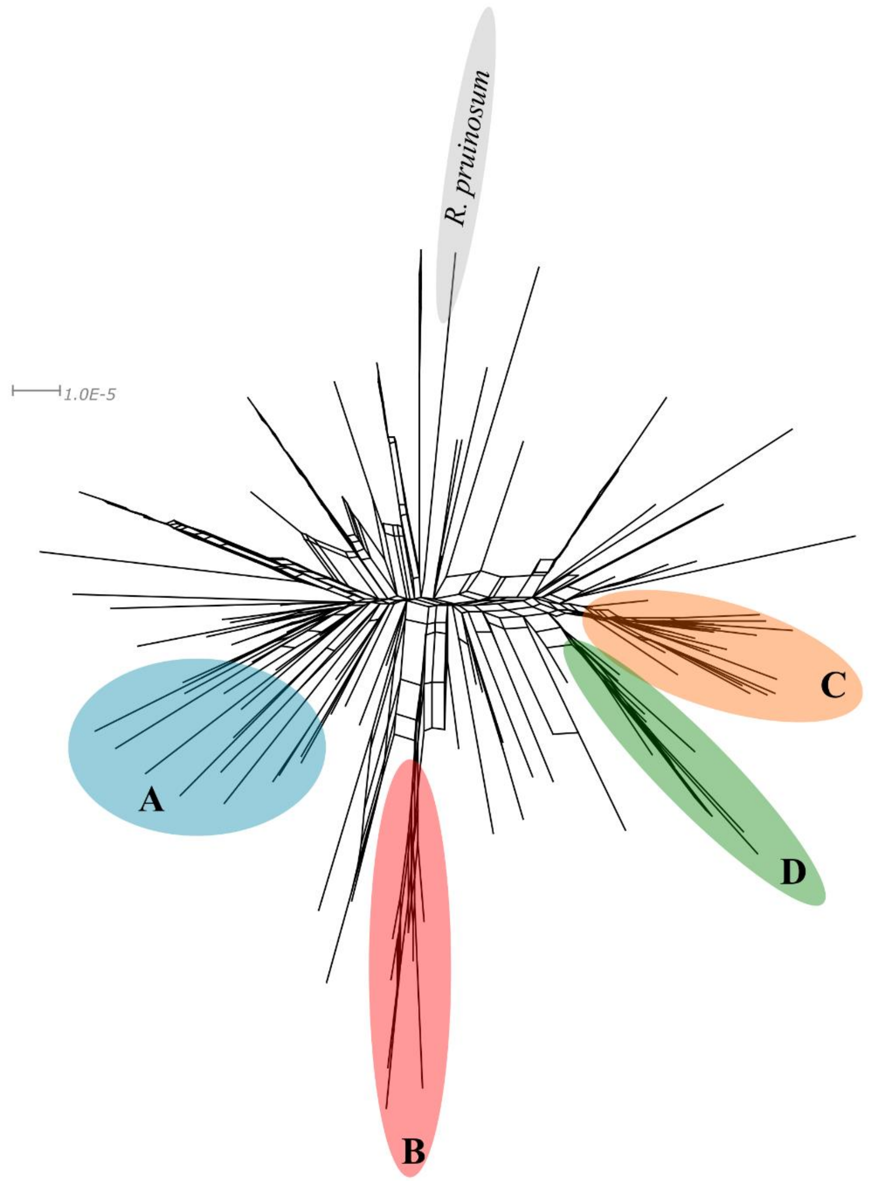


**Figure S4** Phylogenetic network of *Racomitrium lanuginosum* based on uncorrected p-distances of 2718 concatenated loci. The selected loci are shared by at least 20% of 230 samples (-*R* 20, species dataset). The outgroup is represented by *R pruinosum*. Colour codes are analogous to genetic groups in the coancestry matrix. Samples of the tundra are not highlighted.

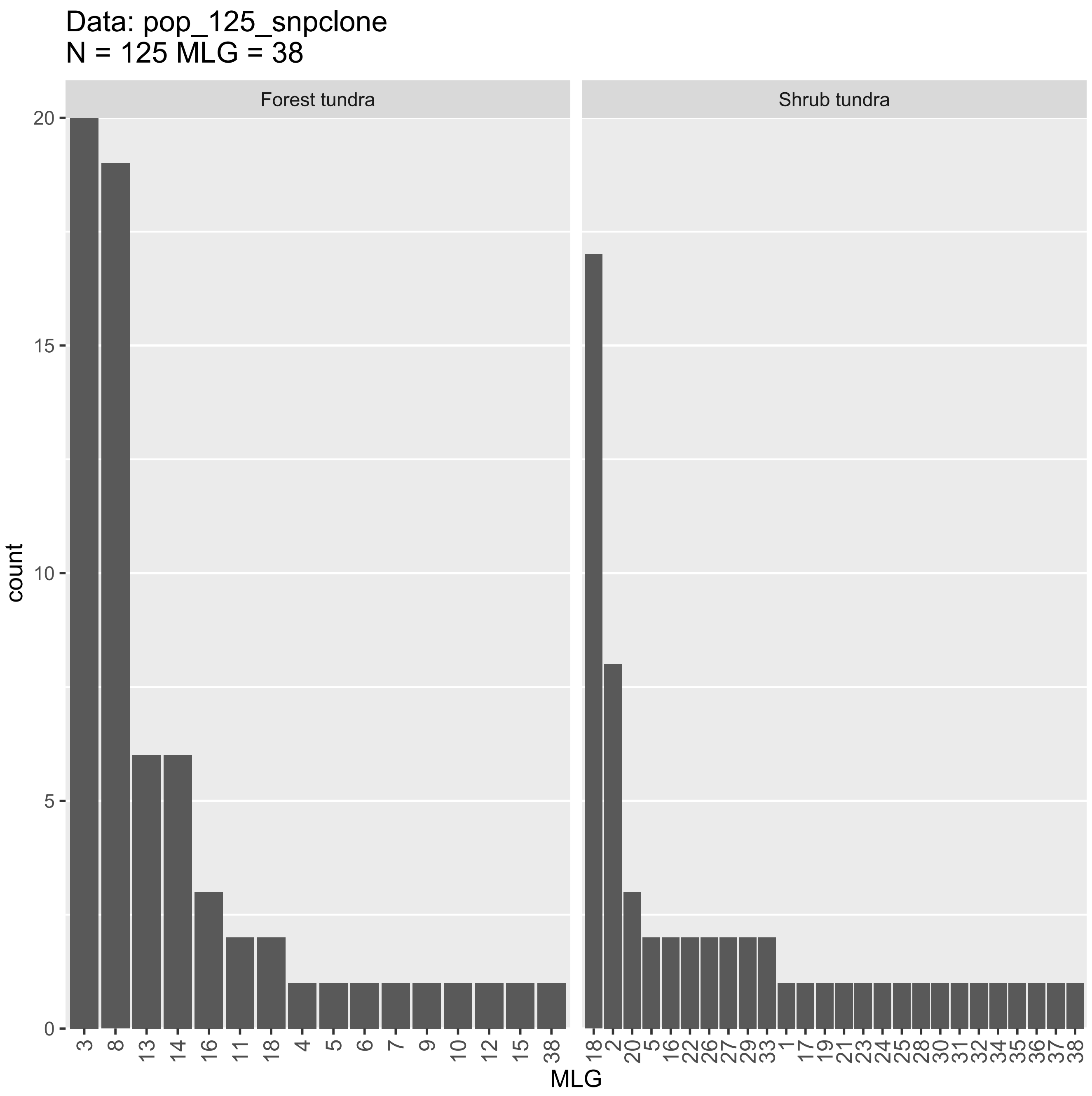


**Figure S5** Multilocus genotype (MLG) counts of *Racomitrium lanuginosum* tundra samples (fine and population scale) per habitat type. The MLGs were inferred from 611 SNPs of the tundra samples (n=125; -*r* 80). The analyses recovered 38 MLGs. Note that the number of samples per tundra type is different.

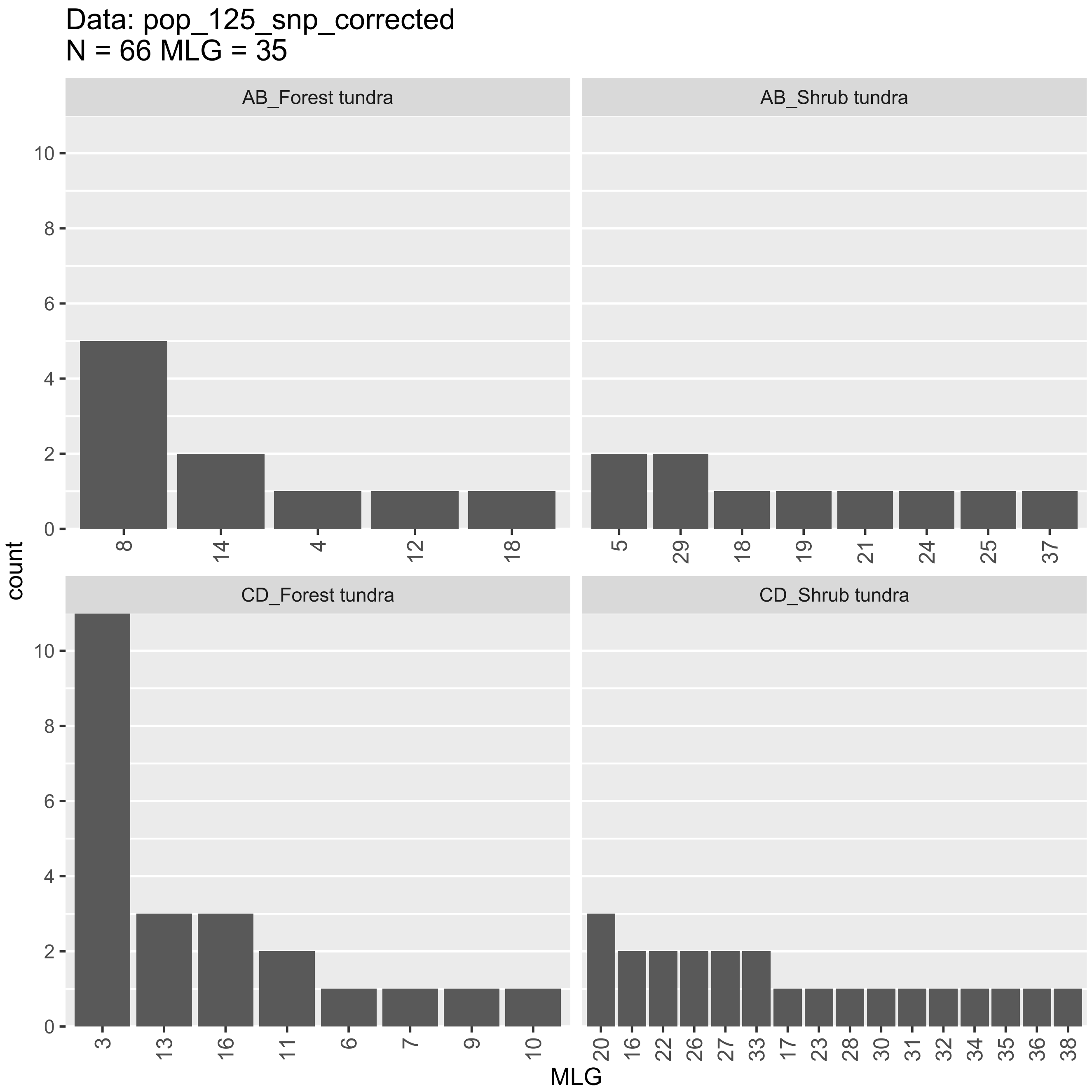


**Figure S6** Multilocus genotype (MLG) counts of *Racomitrium lanuginosum* tundra samples (fine and population scale) per genetic group and habitat type with an even number of samples. The MLGs were inferred from 611 SNPs of the tundra samples (n=66; *-r* 80). Note that the number of samples per genetic group is different

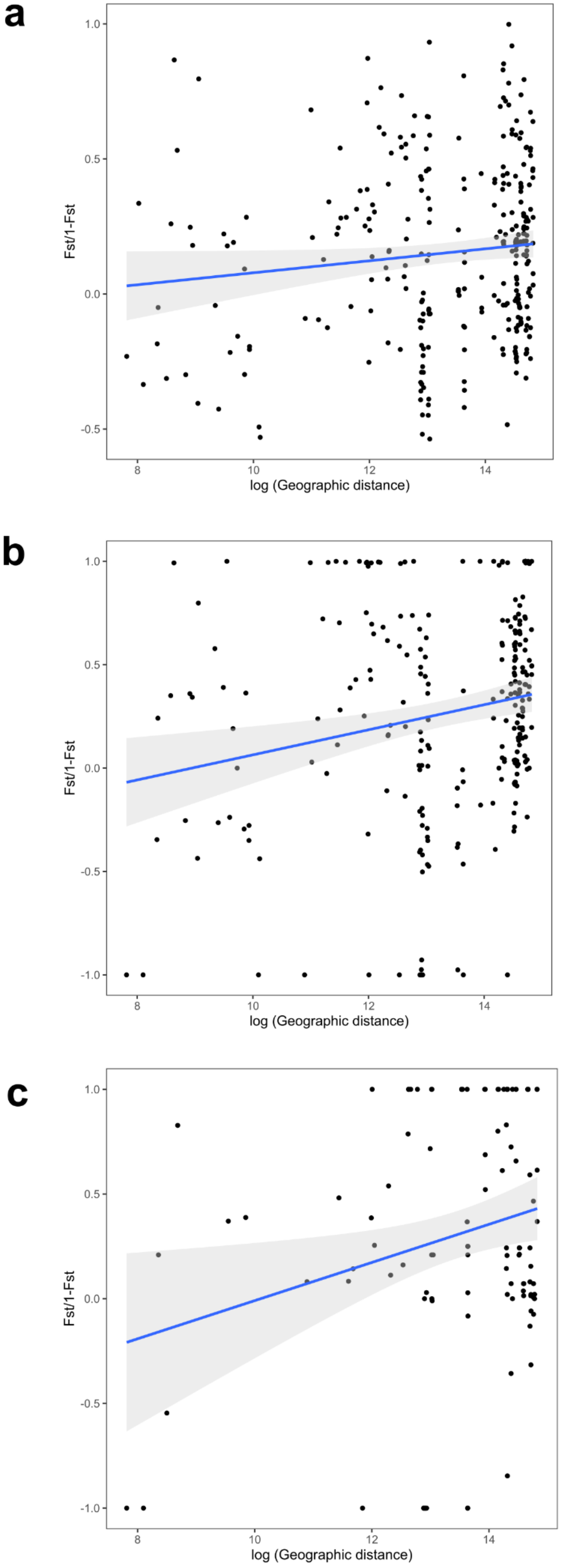
a
b
c
Fst/1-Fst
log (Geographic distance)
1.0
0.5
0.0
-0.5
-1.0
8
10
12
14

**Figure S7.** Isolation by distance relationships of *Racomitrium lanuginosum* samples from the forest tundra and the shrub tundra. **a** Analyses of all samples without genetic group partitioning. Each point indicates pairwise comparisons between plots. The x-axis represents plot-based $F_{ST}$, and the y-axis is the log of the geographic distance (Euclidean) between plots. The blue line indicates the best linear correlation, and the gray area is the 95% confidence interval. There is no strong isolation pattern in the tundra populations when combining both genetic groups ($R^2$= 0.0129, Mantel R statistic= 0.1138, p-value=0.011). **b** Analyses of the genetic group AB. The graph elements represent the same as mentioned before. The genetic group AB has no strong isolation pattern ($R^2$=0.0395, Mantel R statistic=0.1987, p-value=0.001). **c** Analyses of the genetic group CD. The graph elements represent the same as mentioned before. Genetic group CD has no strong isolation pattern ($R^2$=0.0538, Mantel R statistic=0.232, p-value=0.006).

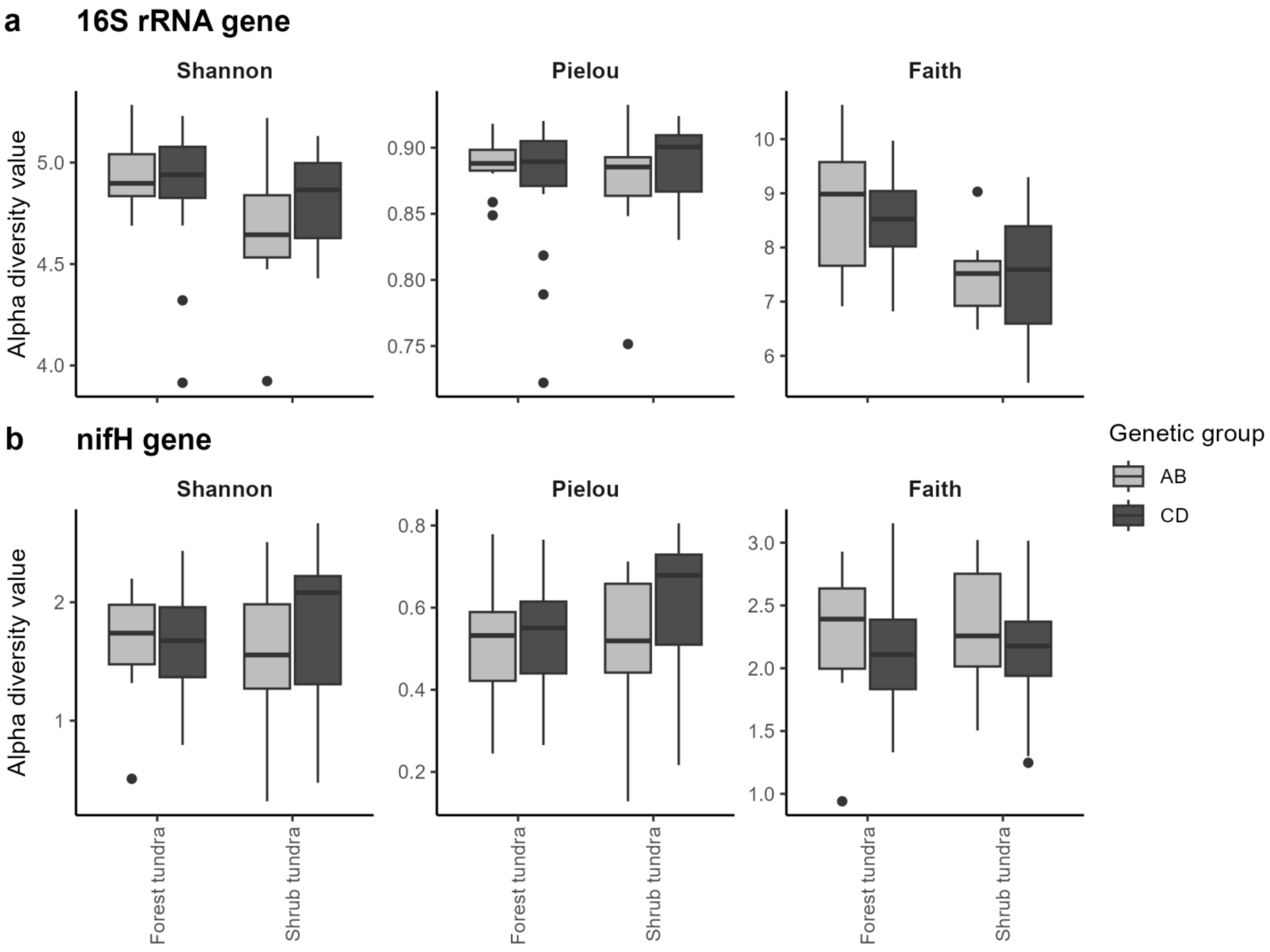


**Figure S8.** Microbial alpha diversity associated with *Racomitrium lanuginosum* genetic groups in the forest tundra ecotone. Shannon, Pielou and Faith's phylogenetic diversity are presented for each habitat and genetic group (in colors). The box represents the first quartile, median and third quartile, with whiskers indicating the maximum and minimum values and outliers as black points for each habitat. **a** Alpha diversity indexes of bacterial communities based on the 16S rRNA gene; **b** Alpha diversity index of diazotrophic community based on the *nifH* gene.

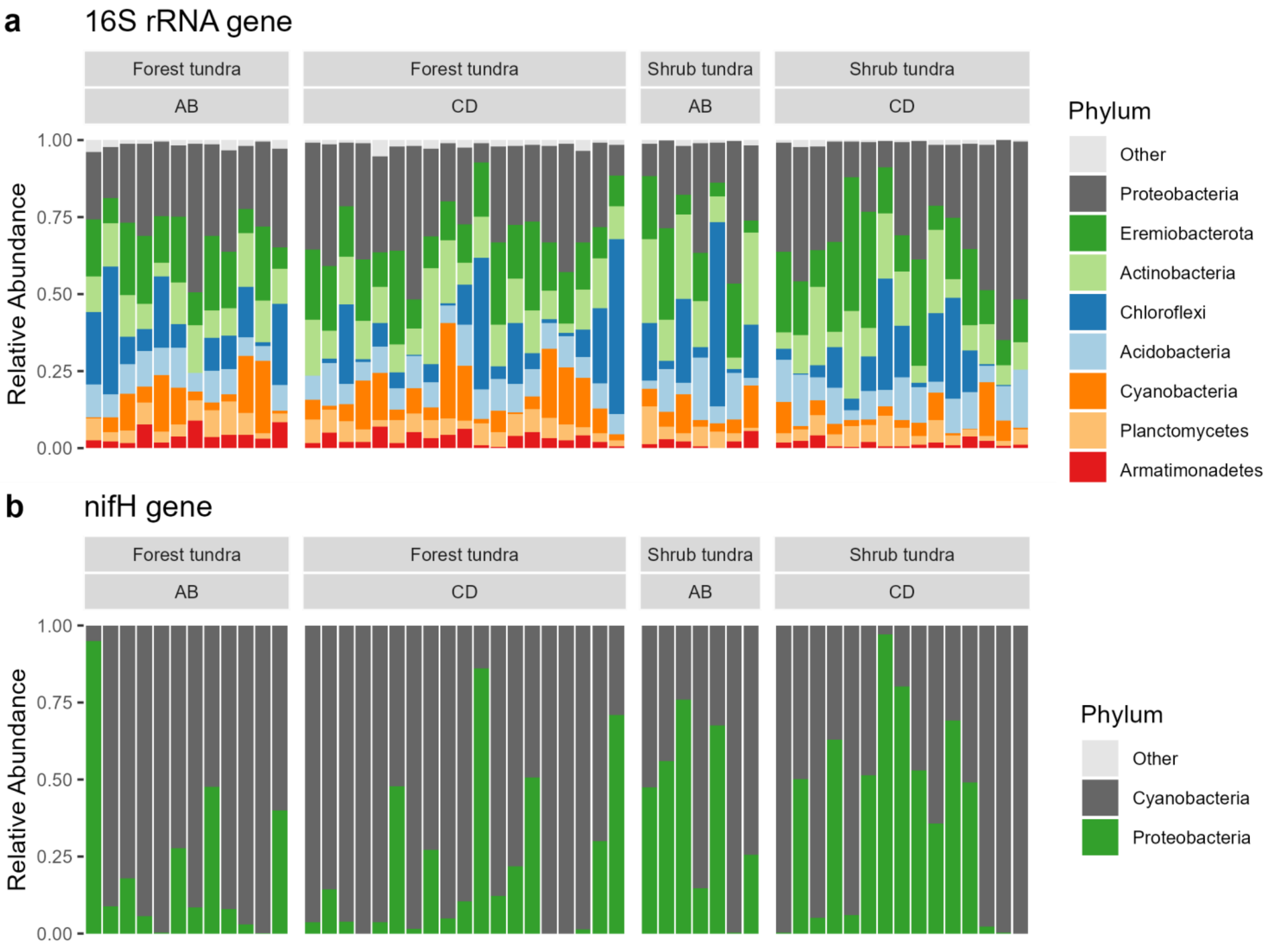


**Figure S9.** Microbial profiles at phylum level associated with *Racomitrium lanuginosum* genetic groups. **a** Relative abundance of bacterial phyla based on the 16S rRNA gene per habitat and genetic group. The most abundant phyla are Proteobacteria, Eremiobacterota (WPS-2) and Actinobacteria. **b** Relative abundance of diazotrophic phyla based on the *nifH* gene per habitat and per genetic group. Cyanobacteria is the most abundant group.

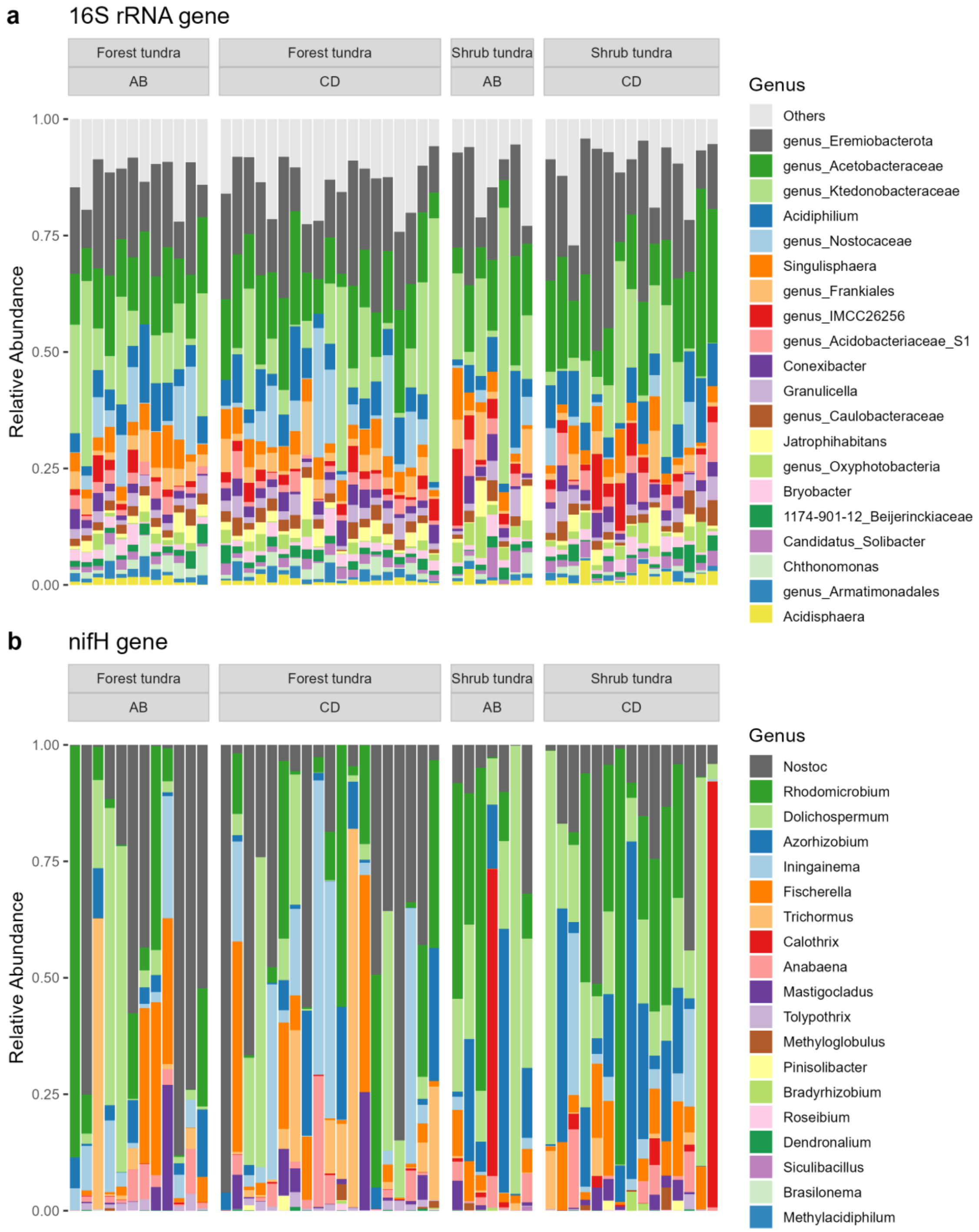


**Figure S10.** Microbial profiles at genus level associated with *Racomitrium lanuginosum* genetic groups. **a** Relative abundance of bacterial genera based on the 16S rRNA gene per habitat and genetic group. **b** Relative abundance of diazotrophic genera based on the *nifH* gene per habitat and per genetic group.

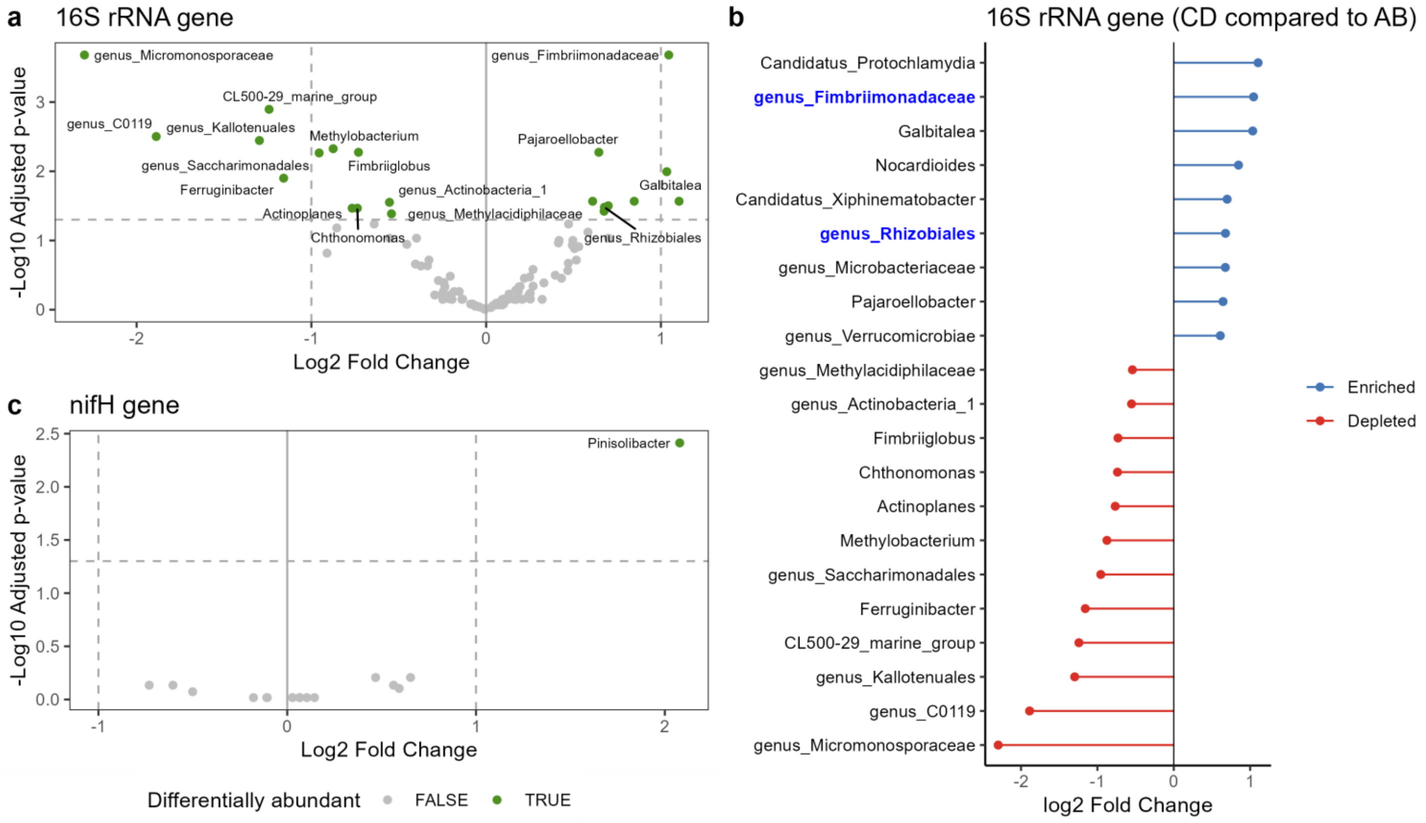


**Figure S11.** Differentially abundant microbial genera associated with *Racomitrium lanuginosum* genetic groups. **a** Volcano plot of adjusted p-values for bacterial abundance in ANCOMBC2 and log2 fold changes (log10 scale) showing differential abundant bacteria in green. **b** Differentially abundant bacterial genera in the genetic group CD compared to group AB based on log2 fold change results. Names in bold blue indicate bacterial genera that passed sensitivity analyses in ANCOMBC2. Only two genera assigned to Fimbriimonadaceae and Rhizobiales were statistically enriched in *R. lanuginosum* samples of genetic group CD compared to AB. **c** Volcano plot of adjusted p-values for diazotrophic abundance in ANCOMBC2 and log2 fold changes (log10 scale) showing differential abundant bacteria in green. Only Pinisolibacter was detected but did not pass the sensitivity test.
